# Structural basis for the tryptophan sensitivity of TnaC-mediated ribosome stalling

**DOI:** 10.1101/2021.03.31.437805

**Authors:** Anne-Xander van der Stel, Emily R. Gordon, Arnab Sengupta, Allyson K. Martínez, Dorota Klepacki, Thomas N. Perry, Alba Herrero del Valle, Nora Vazquez-Laslop, Matthew S. Sachs, Luis R. Cruz-Vera, C. Axel Innis

## Abstract

Free L-tryptophan (L-Trp) induces the expression of the *Escherichia coli* tryptophanase operon, leading to the production of indole from L-Trp. Tryptophanase operon expression is controlled via a mechanism involving the tryptophan-dependent stalling of ribosomes engaged in translation of *tnaC*, a leader sequence upstream of *tnaA* that encodes a 24-residue peptide functioning as a sensor for L-Trp. Although extensive biochemical characterization has revealed the elements of the TnaC peptide and the ribosome that are responsible for translational arrest, the molecular mechanism underlying the recognition and response to L-Trp by the TnaC-ribosome complex remains unknown. Here, we use a combined biochemical and structural approach to characterize a variant of TnaC (R23F) in which stalling by L-Trp is enhanced because of reduced cleavage of TnaC(R23F)-peptidyl-tRNA. In contrast to previous data originated from lower resolution structural studies, we show that the TnaC–ribosome complex captures a single L-Trp molecule to undergo tryptophan-dependent termination arrest and that nascent TnaC prevents the catalytic GGQ loop of release factor 2 from adopting an active conformation at the peptidyl transferase center. In addition, we show that the conformation of the L-Trp binding site is not altered by the R23F mutation. This leads us to propose a model in which rates of TnaC-peptidyl-tRNA cleavage by release factor and binding of the L-Trp ligand to the translating ribosome determine the tryptophan sensitivity of the wild-type and mutant TnaC variants. Thus, our study reveals a strategy whereby a nascent peptide assists the bacterial ribosome in sensing a small metabolite.

## INTRODUCTION

The *tnaCAB* operon of *Escherichia coli* (*E. coli*) contains two structural genes required for the catabolism of L-tryptophan (L-Trp), *tnaA* and *tnaB*, encoding tryptophanase and a tryptophan-specific permease, respectively^1,2^. Tryptophanase catalyzes the breakdown of L-Trp into pyruvate, ammonia and indole^3^, a volatile signaling molecule that affects numerous biological processes within polymicrobial communities^4^. Transcription initiation of *tnaCAB* is controlled by catabolite repression^5^, while the expression of *tnaA* and *tnaB* is regulated via a transcription attenuation mechanism resulting from the tryptophan-dependent stalling of a ribosome producing TnaC, a 24-amino acid leader peptide (Fig. 1a)^6^. When L-Trp levels are low, ribosomes translating *tnaC* dissociate from the mRNA upon reaching the UGA stop codon, leaving an exposed Rho utilization (*rut*) site immediately downstream of *tnaC* that allows recruitment of the Rho transcription termination factor. Rho can then interact with paused RNA polymerase, promoting transcription termination before *tnaA* and *tnaB* are transcribed^6–8^. In the presence of inducing L-Trp levels, translation termination is inhibited when Pro24 of TnaC is in the ribosomal P-site and the stop codon is in the A-site, causing the ribosome to stall. This blocks Rho’s access to the *rut* site, allowing transcription of *tnaA* and *tnaB* to proceed^1,6,7^.

**Figure 1.**
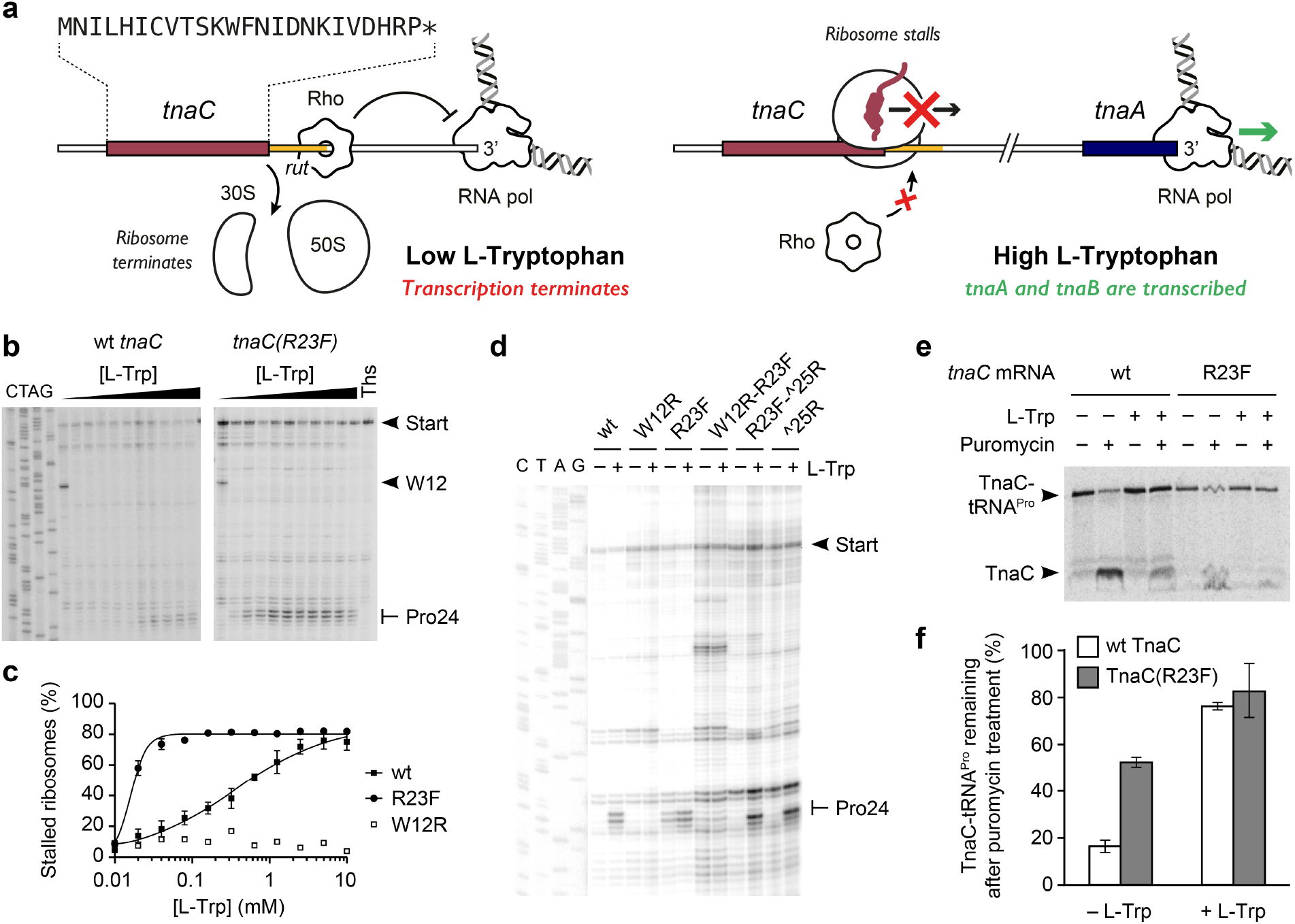
Biochemical characterization of a TnaC(R23F) variant. **a**, Graphical representation of the mechanism of gene regulation by TnaC. A high concentration of L-Trp stalls the ribosome during translation of *tnaC*, obscuring the *rut* site and allowing unhindered transcription of the complete *tnaCAB* operon. **b**, Toeprinting assay of *tnaC* wt and R23F over a range of L-Trp concentrations. Thiostrepton (Ths), an inhibitor of translation elongation, was used as a control for the detection of arrested ribosomes. **c**, Plot of accumulation of stalled ribosomes as a function of L-Trp concentration. Each data point represents the average of three independent experiments. **d**, Toeprinting assay of *tnaC* variants featuring arginine (wt) or phenylalanine (R23F) codons at position 23, an arginine codon at position 25 (^25R) or the double mutation R23F/^25R, as well as W12R or W12R-R23F mutations. Reactions were performed with 0.3 mM (–) or 4 mM (+) L-Trp. **e**, Radiogram of resolved products of puromycin-induced cleavage of [^35^S]-methionine labeled TnaC-tRNA molecules. The complexes were incubated with (+) or without (–) 4 mM L-Trp and later incubated with (+) or without (–) 1 mM puromycin. Arrows indicate the position of the [^35^S]-methionine TnaC-tRNA^Pro^ and TnaC-puromycin (TnaC) products. **f**, Bar plot indicating the relative remaining amount of TnaC-tRNA^Pro^ in the wild type and R23F mutant complexes after treatment with puromycin. These results are indicative of three independent experiments.

The tryptophan-mediated stalling of ribosomes translating *tnaC* requires elements from both the nascent peptide and the ribosome. Changes to four critical TnaC residues (Trp12, Asp16, Ile19 and Pro24)^9–14^ or to specific elements within the 23S ribosomal RNA (rRNA) or ribosomal proteins uL4 and uL22 belonging to the large ribosomal subunit^15–18^ abolish TnaC-mediated stalling in response to L-Trp. Low (5.8 Å)^19^ to medium (3.2–3.8 Å)^20,21^ resolution cryogenic electron microscopy (cryo-EM) analyses of TnaC–ribosome complexes have suggested possible interactions between these functional residues of TnaC and elements constituting the peptidyl transferase center (PTC) or nascent polypeptide exit tunnel of the ribosome, but have either failed to identify the bound L-Trp^19^ or hinted that the TnaC–ribosome complex may interact with one^21^ or two distinct L-Trp molecules^20^. Consequently, a detailed understanding of the mechanism by which a TnaC–ribosome complex senses L-Trp is still missing.

Here, we use a combined forward genetic selection and screening approach to identify a TnaC variant that functions in the same manner as wild-type TnaC, albeit with greatly enhanced sensitivity for L-Trp. Using single-particle cryo-EM, we obtained a high-resolution structure (2.4 Å) that shows that this variant adopts a compact structure within the ribosome that is virtually indistinguishable from that of wild-type TnaC, but differs considerably from the one observed in an earlier medium-resolution reconstruction of the TnaC–ribosome complex^20^. In particular, our structural and biochemical analyses reveal the molecular details leading to the capture of a single L-Trp molecule by the TnaC–ribosome complex. This allows us to propose a model for tryptophan-dependent translational arrest in which the relatively low rate of TnaC peptidyl-tRNA hydrolysis by release factor 2 (RF2) gives sufficient time for free L-Trp to bind to the ribosome and stabilize a TnaC conformation that silences the PTC, resulting in ribosome stalling.

## RESULTS

### A single R23F mutation increases the tryptophan sensitivity of TnaC

We hypothesized that intragenic mutations that suppress a loss-of-function *tnaC* variant would reveal elements that enhance translational arrest in a wild-type background, possibly even voiding the need for L-Trp as an inducer. Such elements might include TnaC residues that modulate the basal response observed *in vivo*, which is thought to occur when a ribosome approaching the stop codon of *tnaC* undergoes a tryptophan-independent pause, preventing Rho from binding to the *rut* site and reducing its ability to abort transcription^14^. To generate a loss-of-function variant, we targeted Asp16, a functionally important residue^14,15^ that has been suggested not to interact directly with the L-Trp ligand^20^. Nine different amino acid substitutions at position 16 of TnaC are known to abolish tryptophan-dependent ribosome stalling, including the conservative D16E mutation^12,15^. Using a combined forward genetic selection and screening approach, we identified second site mutations at position 10 (S10P) or 23 (R23H or R23S) that suppress the non-functional D16E *E. coli* TnaC variant (Supplementary Notes, Supplementary Table 1). As expected, incorporation of the S10P or R23H single mutations into the wild-type *tnaC* gene resulted in a ∼20-fold increase of the basal level of expression of a *tnaA-lacZ* fusion reporter gene (Supplementary Table 1). Substitutions at position 23 are particularly interesting in light of recent data showing that Arg23 helps maintain a weak wild-type basal response *in vivo*^14^. Furthermore, *tnaC* homologues in different bacterial species where ribosome stalling occurs at elongation rather than translation termination encode Phe at the position corresponding to the 23^rd^ codon of the *E. coli tn*aC gene^22,23^. It is possible that in these *tnaC* homologues a phenylalanine residue may have been selected over arginine to reduce the rate of amino acid incorporation and bring it closer to that of peptidyl-tRNA hydrolysis^24^. We therefore investigated the ability of the TnaC(R23F) single mutant to cause ribosome stalling, anticipating that this variant would have a reduced dependency on L-Trp or even be capable of undergoing constitutive translation termination arrest.

To compare the ability of wild-type TnaC and TnaC(R23F) to stall the ribosome *in vitro* in response to different concentrations of L-Trp, we performed toeprinting assays to monitor the position of ribosomes on transcripts encoding these two variants (Fig. 1b). For wild-type TnaC, the concentration of L-Trp necessary to obtain 50% of the maximum accumulation of arrested ribosomes on codon 24 of *tnaC* ([L-Trp]_0.5_) was 0.3 mM, with a maximum accumulation observed at concentrations greater than 5 mM (Fig. 1c). In contrast, [L-Trp]_0.5_ for TnaC(R23F) was nearly 30-fold lower and maximum ribosome accumulation was achieved at ligand concentrations of only ∼0.1 mM (Fig. 1c). These data suggest that nascent TnaC(R23F) significantly increases the ribosome sensitivity to L-Trp to stall at the termination site of *tnaC*. Incorporating the W12R mutation, which is known to abolish TnaC-mediated ribosome stalling (Fig. 1c)^10^, into the TnaC(R23F) variant (W12R-R23F), abrogates ribosome stalling at the end of the *tnaC* ORF (Fig. 1d), suggesting that the arrest caused by the wild-type and R23F TnaC peptides operates through similar mechanisms.

Because the identity of the amino acid residue at position 23 differs between natural TnaC sequences that inhibit translation termination or elongation^22^, we asked whether the TnaC(R23F) variant could also direct elongation arrest at low L-Trp concentrations. Toeprinting analysis comparing wild-type and R23F *tnaC* with similar variants where the stop codon at position 25 was substituted by an arginine codon (^25R and R23F–^25R), indicated that the R23F substitution did not lead to elongation arrest at low L-Trp concentrations (0.3 mM) (Fig. 1d). However, arrest of translation elongation was still observed in these constructs at high L-Trp concentration (4 mM) (Fig. 1d). Consistent with the results of toeprinting analysis, *in vitro* translation of the template encoding TnaC(R23F), unlike that with the R23F–^25R construct, results in accumulation of TnaC–tRNA^Pro^ at low L-Trp concentrations (Supplementary Fig. 1). These results indicate that the enhanced sensitivity to L-Trp mediated by the TnaC(R23F) peptide to arrest translation operates during termination at codon 25, but not during elongation at codon 25 of the *tnaC* ORF.

We next sought to determine whether ribosome stalling induced by the R23F mutant was dependent on small amounts of L-Trp present in the translation reaction or if the translational arrest was truly constitutive. To test this, we determined the ability of the ribosome to transfer the different nascent TnaC peptide variants from TnaC–tRNA^Pro^ to puromycin, an antibiotic that mimics aminoacyl tRNA in the ribosomal A site, under conditions where soluble L-Trp can be completely depleted. Therefore, we isolated ribosome complexes containing wild-type or R23F TnaC–tRNA^Pro^ and challenged them with puromycin in the presence or absence of L-Trp^16^ (Fig. 1e). In the presence of L-Trp, the extent of puromycin protection was similar for the wild-type and R23F variants (75% and 82% respectively) (Fig. 1f). In the absence of L-Trp, even though the complexes carrying the TnaC(R23F) peptide variant were three times more resistant to puromycin than those with wild-type TnaC, their sensitivity to puromycin cleavage remained as high as 50% (Fig. 1f). These data indicate that, while the R23F mutation somehow hinders the transfer of the peptide to puromycin, it does not produce a truly constitutive arrest peptide. Altogether, these data show that the TnaC(R23F) peptide variant undergoes strong termination arrest that is hypersensitive to L-Trp, but that L-Trp is nevertheless necessary to ensure robust TnaC(R23F)–ribosome stalling.

### TnaC(R23F) and wild-type TnaC adopt the same compact structure

In order to understand how the R23F mutation leads to increased tryptophan sensitivity, we prepared *E. coli* 70S ribosomes stalled during translation of *tnaC(R23F)* in the presence of L-Trp and analyzed their structure using cryo-EM (Fig. 2a and Supplementary Figs. 2 and 3). A major subpopulation (∼53% of particles) corresponding to ribosomal complexes with TnaC(R23F)– tRNA^Pro^ in the ribosomal P-site could be identified, resulting in a structure with an overall resolution of 2.4 Å, referred to as TnaC(R23F)–70S. Clear density in the region of the nascent polypeptide exit tunnel between the PTC and the constriction formed by ribosomal proteins uL4 and uL22 allowed us to unambiguously model the backbone and side chains of residues 9– 24 of TnaC(R23F) (Fig. 2b and Fig. 3a). On the other hand, the density corresponding to residues 1–8 was weak to non-existent, consistent with the negligible impact of TnaC N-terminal deletions on ribosome stalling (Supplementary Fig. 4) and previous reports indicating that the nature of the N-terminal residues of TnaC is not important for translational arrest^14^. The peptide conformation we observe does not resemble that in an earlier 3.8 Å resolution cryo-EM reconstruction of the TnaC–ribosome complex^20^, but bears some similarity to the 3.2 Å structure of a titin I27–TnaC fusion^21^. To address this discrepancy, we also determined the structure of a wild-type TnaC–70S complex stalled in the presence of L-Trp at an overall resolution of 2.9 Å (∼10% of particles). Although we observed an overall weaker density for TnaC compared to that of the mutant peptide in the TnaC(R23F)–70S structure (Supplementary Fig. 2), the excellent overlap between our wild-type and R23F peptide densities indicates that the mutation does not alter the conformation of the nascent peptide, with the obvious exception of the side chain of residue 23, which makes different contacts with the ribosome in the wild-type (Arg) and mutant (Phe) structures (Fig. 3b). It is possible that partial peptide occupancy combined with lower resolution might have caused the peptide density to be misinterpreted in the earlier TnaC–ribosome map^20^.

**Figure 2.**
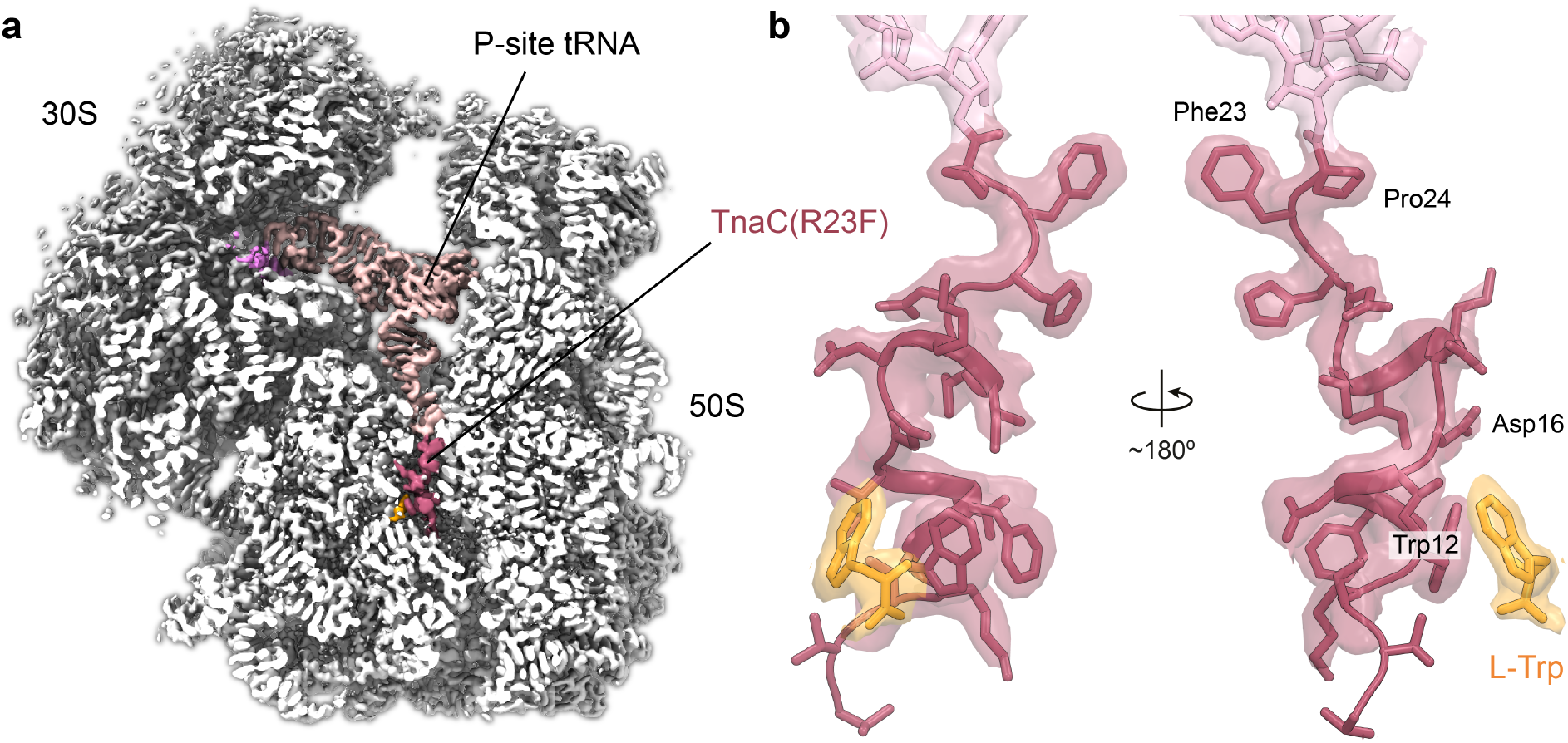
Cryo-EM structure of a TnaC(R23F)–70S complex. **a**, Cross-section of a cryo-EM density map of the TnaC(R23F)-70S complex showing the 70S ribosome (white), the TnaC(R23F) peptide (red) in the nascent polypeptide exit tunnel attached to tRNA^Pro^ (pink) in the ribosomal P-site, and the mRNA (dark pink). A tryptophan molecule can be seen adjacent to the TnaC(R23F) peptide (orange). **b**, Close-ups of the TnaC(R23F) peptide attached to the P-site tRNA and of the L-Trp molecule, showing the modeled structure and the corresponding density. Coloring is as in panel a.

**Figure 3.**
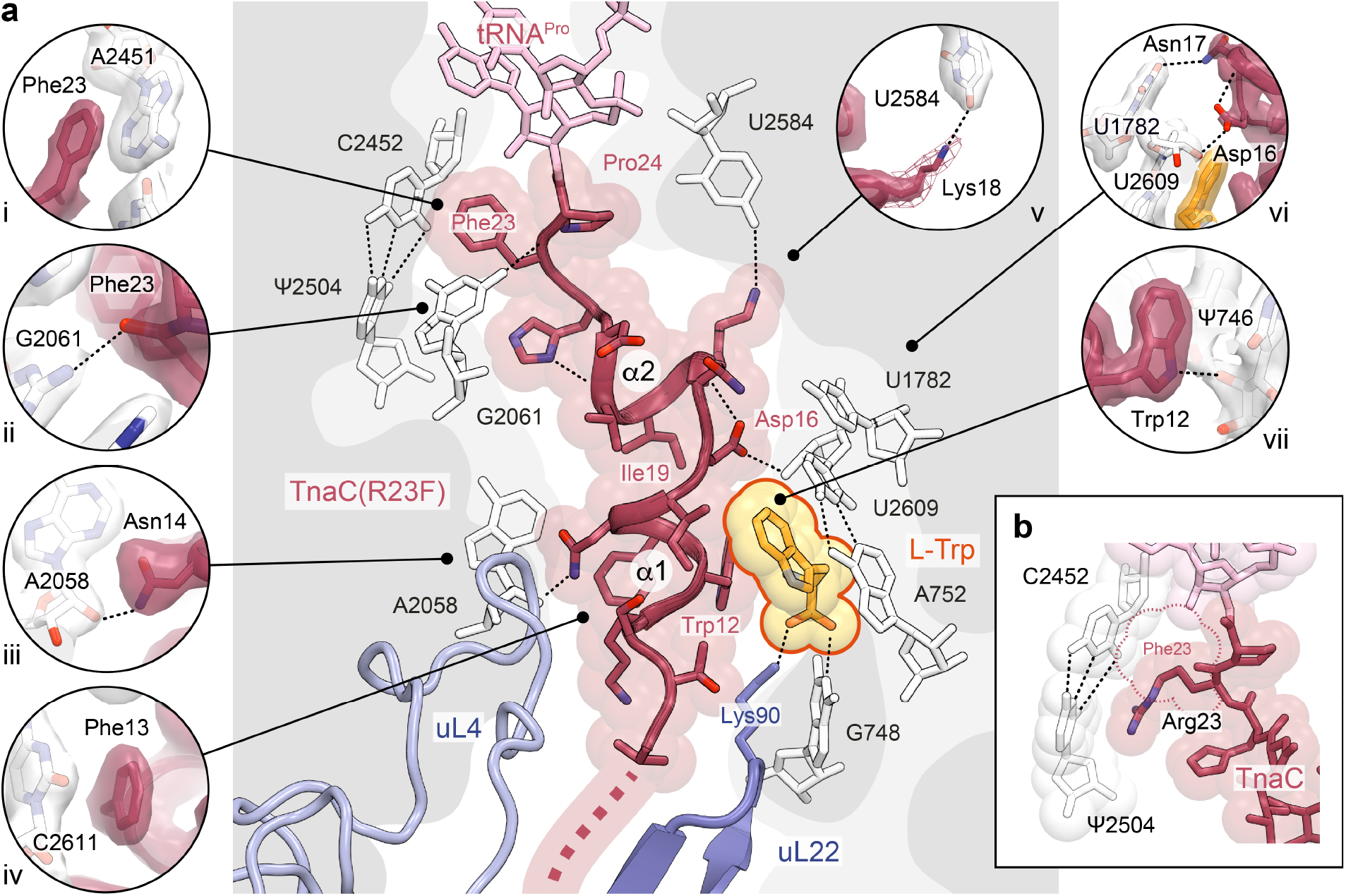
Contacts between TnaC(R23F) and the ribosome. **a**, The central panel shows the TnaC(R23F) nascent chain (red) attached to the P-site tRNA (pink), the L-Trp ligand (orange) and ribosomal proteins uL4 (light blue) and uL22 (periwinkle blue). Prominent contacts with the 23S rRNA (white) are shown in individual panels, including (i) a π-π stacking interaction between the side chain of TnaC residue Phe23 and the base of 23S rRNA residue A2451; (ii) a hydrogen bond between the carbonyl oxygen of TnaC residue Phe23 and the N2 amine of 23S rRNA residue G2061; (iii) a hydrogen bond between the carboxamide nitrogen of TnaC residue Asn14 and the 2’-hydroxyl of 23S rRNA residue A2058; (iv) a π-π stacking interaction between the side chain of TnaC residue Phe13 and the base of 23S rRNA residue C2611; (v) a hydrogen bond between the side chain amine of Lys18 (shown at a lower contour level as a mesh) and the O4 carbonyl of 23S rRNA residue U2584; (vi) hydrogen bonds between the side chain of TnaC residue Asn17 the base of 23S rRNA residue U1782, and between the side chain of TnaC residue Asp16 and the 2’-hydroxyl of 23S rRNA residue U2609; and (vii) a hydrogen bond between the side chain amine of Trp12 and the 2’-OH of 23S rRNA residue Ψ746. **b**, Conformation of wild-type TnaC residue Arg23 and interactions with the neighboring C2452:Ψ2504 base pair of the 23S rRNA in the TnaC–70S structure. The shape of the Phe23 side chain in the TnaC(R23F)–70S structure is shown as a dotted line.

In the stalled TnaC(R23F)–70S and TnaC–70S termination complexes, the C-terminal half of TnaC folds into a compact structure consisting of two short consecutive α-helical segments – α1 (residues 11–15) and α2 (residues 17–21) – connected by a ∼110° hinge centered on the side chain of Ile19 (Fig. 3a). Only medium-sized hydrophobic side chains are tolerated in this position^14,25^, which is consistent with a role for residue 19 in maintaining the relative orientations of α1 and α2. A similar propensity to form short α-helices between the PTC and the tunnel constriction was reported for two other arrest peptides, SpeFL^26^ and VemP^27^. Additional intramolecular interactions that help maintain the structure of TnaC include hydrogen bonds between the carbonyl oxygen of Ile19 and the side chain of His22, and between the side chain carboxyl of Asp16 and the backbone amine of Lys18 (Fig. 3a). Moreover, TnaC establishes numerous contacts with neighboring ribosomal residues. These include a hydrogen bond between the 2’-hydroxyl of 23S rRNA residue Ψ746 and the indole amine of the strictly conserved Trp12 of TnaC (Fig. 3a), a residue whose position in our structure is consistent with crosslinking data^16^ and whose mutation to any other amino acid abolishes stalling^10,11,14^. Similarly, the conserved TnaC residue Asp16 helps anchor the nascent peptide to the ribosome via a hydrogen bond formed between its side chain carboxyl and the 2’-hydroxyl of 23S rRNA residue U2609 (Fig. 3a), in agreement with chemical modification experiments^15^. By acting as a bridge between U2609 and Lys18 of TnaC, Asp16 thus appears to play a key structural role in line with its demonstrated functional importance^22^. The stacking of the phenyl group of TnaC residue Phe13 against the base of 23S rRNA residue C2611 also helps to stabilize the nascent peptide within the tunnel (Fig. 3a), explaining the need for an aromatic side chain at this position^14^. A hydrogen bond between the side chain of TnaC residue Asn14 and the 2’-hydroxyl of 23S rRNA residue A2058 (Fig. 3a) is consistent with the functional role suggested for the latter in the stalling process^25^. These and other interactions with the ribosome help TnaC maintain a compact structure that limits its movement inside the exit tunnel, thereby restricting its ability to maneuver through the uL4-uL22 constriction.

### The TnaC–ribosome complex binds a single L-Trp molecule

In addition to the TnaC peptide, we observed clear density for a single L-Trp molecule near the tunnel constriction (Fig. 2b, Fig. 3a and Fig. 4a)^20^. The indole side chain of L-Trp makes contact with both the ribosome and the nascent peptide, and is wedged between the base pair formed by residues A752 and U2609 of the 23S rRNA, and residues Trp12, Ile15 and Asp16 belonging to helix α1 of TnaC (Fig. 3a and Fig. 4a). Disruption of the A752:U2609 base pair severely impairs TnaC-mediated stalling, while single (A752G) or double (A752G/U2609C) mutations that maintain base pairing preserve the ability of TnaC to undergo tryptophan-dependent stalling^15^. The above-mentioned Asp16–U2609 and Trp12–Ψ746 hydrogen bonds help to position α1 of TnaC with respect to the 23S rRNA, indicating a key role in defining the correct geometry of the binding pocket that further confirms their functional importance. While residues from TnaC and the 23S rRNA act in concert to bind the indole moiety of L-Trp, the ligand backbone is recognized exclusively by the ribosome. First, the α-carboxyl group of L-Trp makes a hydrogen bond with the Watson-Crick edge of the 23S rRNA G748 base and a salt bridge with the side chain amine of residue Lys90 from ribosomal protein uL22 (Fig. 4). This explains why the deletion of this α-carboxyl group (tryptamine)^28^ or its substitution with hydroxyl (L-tryptophanol)^29^, amide (L-tryptamide)^28^ or methyl ester (L-tryptophan methyl ester)^28^ moieties lacking a negative charge result in a non-functional ligand, and why mutations at position 90 of uL22 (K90H and K90W) eliminate tryptophan-dependent stalling^16^. Second, the α-amino group of L-Trp forms a hydrogen bond with the O4 carbonyl oxygen of Ψ746 (Fig. 4). Not surprisingly, a tryptophan analogue lacking an α-amino group (indole-3-propionic acid) fails to induce TnaC-mediated arrest^28^, but addition of a glycyl moiety to the α-amine of L-Trp yields a functional ligand (glycyl-L-tryptophan) capable of inhibiting TnaC–tRNA^Pro^ hydrolysis at the PTC^28^, an observation that is consistent with the dimensions of the vacant space adjacent to the α-amino group in our structures. While the hydrogen bonding pattern imposed by residues Ψ746, G748 and Lys90 precludes interactions with the backbone of D-tryptophan^28^, side chain selectivity is ensured by the shape, size and chemical properties of the binding pocket. The π-π interaction observed between the L-Trp side chain and the A752:U2609 base pair mirrors that between this base pair and the alkyl-aryl moiety of the ketolide antibiotic telithromycin^30,31^, indicating that this region of the exit tunnel favors interactions with aromatic ligands via π-stacking. Moreover, the ∼3.5 Å gap separating Trp12 of TnaC from base pair A752:U2609 (Fig. 4a) may act as a molecular sieve to allow aromatic side chains into the binding pocket while excluding the thicker side chains of all but the smallest amino acids. This idea is consistent with the inability of an L-Trp analog with a non-planar side chain (L-7-aza-tryptophan) to induce arrest^28^. Aromatic side chains that succeed in entering this cavity would engage in π-stacking with the A752:U2609 base pair, but the smaller side chains of histidine, phenylalanine and tyrosine would likely result in a looser fit, explaining their inability to trigger arrest^27^. The importance of van der Waals interactions for optimal side chain accommodation is illustrated by the effect of various methyl groups added to the L-Trp side chain, which can have a deleterious or neutral impact on TnaC-mediated PTC shutdown depending on whether they are added at the C4/C6 or N1/C5 positions, respectively^28^. Surprisingly, the indole amine of L-Trp is not involved in hydrogen bonding and can be methylated without affecting its ability to cause arrest^28^, even though it could potentially be used to discriminate free tryptophan from phenylalanine or tyrosine. Not relying on this group for specificity could in fact prevent the formation of a hydrogen bond leading to the undesirable binding of free histidine or indole, the product of tryptophan degradation that accumulates upon expression of tryptophanase. In short, our data provide a structural explanation for the ability of TnaC to exclude indole and all amino acids other than L-Trp from the ligand-binding pocket it helps create inside the ribosome.

**Figure 4.**
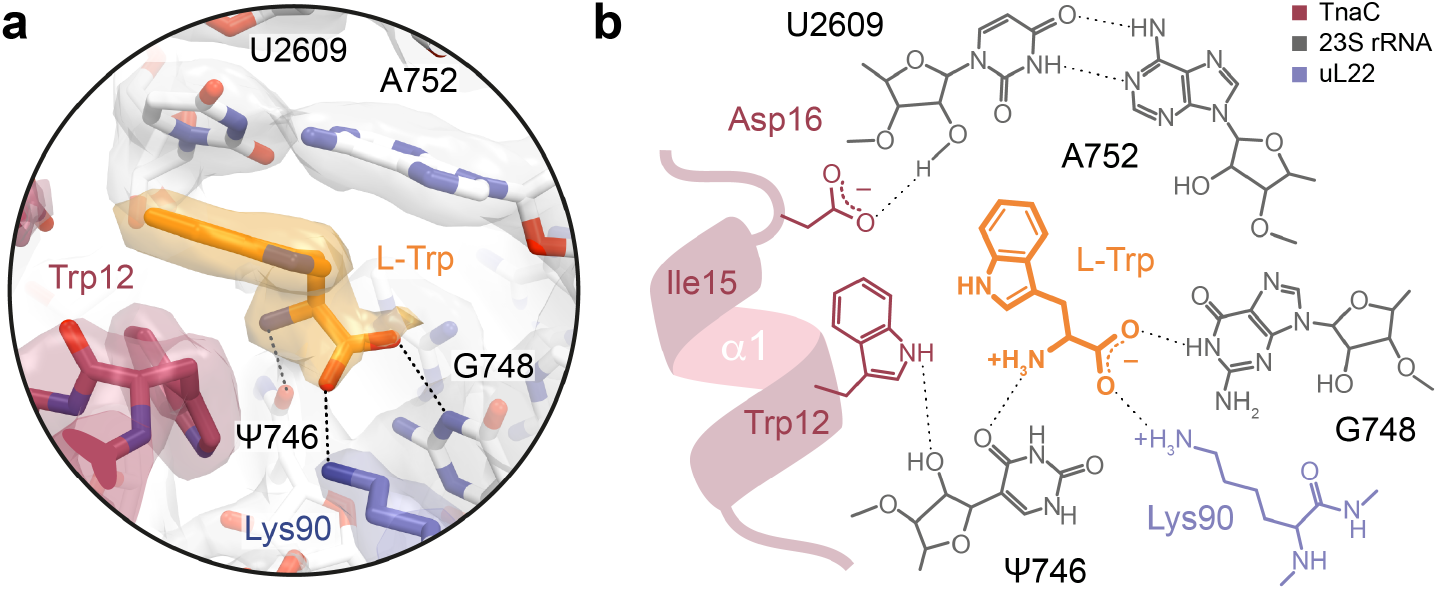
The TnaC–ribosome complex captures a single L-Trp molecule. **a**, Close-up of the L-Trp binding site, showing the models and corresponding densities for the TnaC(R23F) peptide (red), L-Trp (orange), 23S rRNA (white) and ribosomal protein uL22 (periwinkle). The L-Trp ligand is wedged between residue Trp12 of TnaC and the 23S rRNA base pair A752:U2609, with which it makes a π-π stacking interaction. **b**, Chemical diagram showing the binding of the L-Trp ligand. The backbone of the L-Trp ligand makes hydrogen bonds with the bases of 23S rRNA residues Ψ746 and G748, and a salt bridge with residue Lys90 of ribosomal protein uL22.

### RF2 binds to the TnaC–ribosome complex but cannot reach the PTC

In addition to the complexes described above, we also observed a low-abundance class corresponding to RF2 bound to TnaC(R23F)–70S (∼15% of particles), which could be refined to yield a structure with an overall resolution of 2.6 Å (Fig. 5a). In the TnaC(R23F)–70S–RF2 complex, domain III of RF2 is shifted relative to the active, canonical termination complex and only very weak, discontinuous density is observed for residues 247–256 of its catalytic GGQ loop (Fig. 5b). Thus, it appears that the presence of TnaC inside the exit tunnel does not prevent domain III from crossing the “accommodation gate” leading to the ribosomal A-site^32^, but stops the GGQ loop from engaging with the PTC. This is consistent with biochemical data showing that the peptidyl-hydrolase activity of RF2 is blocked within the TnaC-ribosome complex^33^. In contrast, interactions between domain II of RF2 and the A-site codon in the mRNA are maintained, indicating that TnaC(R23F)-mediated ribosome stalling does not interfere with RF2 recognition of the UGA stop codon, in agreement with earlier observations^8,16,28^ (Supplementary Fig. 5). Although we could not detect a class containing RF2 for the wild-type TnaC–70S complex, pull-down experiments with biotinylated *tnaC* mRNA demonstrated the existence of a stable TnaC–70S–RF2 complex (Fig. 5c). Introducing the S205P mutation into RF2, which is known to disrupt the interaction between domain II and the A-site codon^34^, completely suppressed the TnaC-70S-dependent pull-down of RF2. In contrast, the Q252A mutation in the GGQ loop that reduces the RF2-dependent rate of hydrolysis of peptidyl-tRNA without interfering with the recognition of the stop codon^35^ had no effect on the interaction between RF2 and the stalled complex (Fig. 5c). Thus, while RF2 can associate with the stalled TnaC– ribosome complex by interacting with the A-site codon, its GGQ loop is unable to fold into a productive conformation and catalyze the hydrolysis of peptidyl-tRNA.

**Figure 5.**
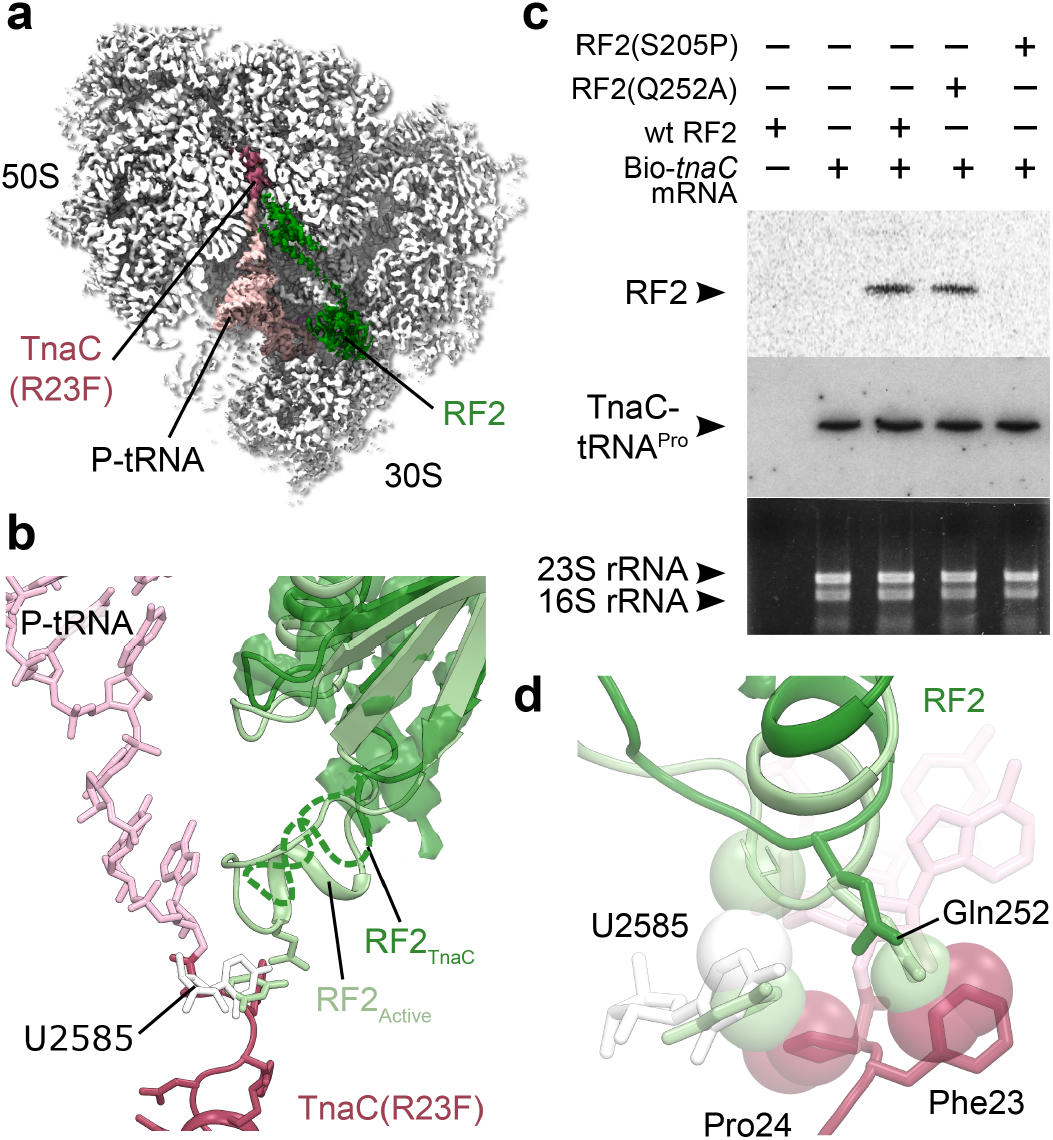
RF2 binds to TnaC(R23F)-70S but does not accommodate into the PTC. **a**, Cross section of the TnaC(R23F)–70S–RF2 density map showing the 70S ribosome (white), the P-site tRNA (pink), the TnaC nascent chain (red) and RF2 (green). **b**, RF2 bound to the TnaC–ribosome complex (dark green) fails to accommodate into the PTC as indicated by the lack of density for the GGQ loop. Active RF2 (light green) is shown for comparison (PDB 4V67^63^). The RF2 bound to the 70S-TnaC ribosome is shifted ∼2Å away from the PTC. Besides, the RF2 residues 247 to 256, including the crucial GGQ motif, show a very weak, non-continuous density (dark green surface), indicating a disordered, inactive conformation. **c**, Pull-down experiment showing RF2 can bind to the 70S-TnaC-wt complex. RF2-wt as well as the Q252A mutant are retained by TnaC-stalled ribosomes, in contrast to the S205P mutant, which is critical for the UGA codon recognition. **d**, Close-up of the PTC area. The conformation of RF2 Q252 in the active conformation is clashing with the F23 residue of the TnaC nascent chain. Moreover, the U2585 is pushed back by P24 of the nascent chain (white sticks) compared to the conformation in the active RF2 complex (light green sticks) (PDB 4V67) and clashes with the RF2 in the active conformation.

Although previous structures of stalled TnaC–ribosome complexes did not capture the RF2-bound state, they showed that 23S rRNA residues A2602 and U2585 adopt conformations that are incompatible with the binding of RF2^19,20^. If we first consider the rather weak cryo-EM density for A2602 in all of the structures presented here, it seems likely that the conformation we observe represents the most stable among a number of possible conformational states. This, in turn, suggests that A2602 could move aside to allow RF2 binding, making it an unlikely candidate for PTC silencing. On the other hand, the TnaC(R23F)–70S and TnaC(R23F)–70S–RF2 structures clearly show that 23S rRNA residue U2585 and TnaC residue Phe23 would clash with Gly251 and Gln252 of the RF2 GGQ loop, respectively, likely preventing RF2 from accommodating fully into the PTC (Fig. 5d). In the TnaC–70S structure, Arg23 of TnaC would also clash with Gln252 in the position it occupies in the canonical termination complex, albeit less severely than Phe23 of TnaC(R23F). It is therefore possible that RF2 would primarily need to surmount a clash with U2585, which would itself be caused by U2585 being pushed away from its normal RF2-bound conformation by Pro24 of TnaC. Mutating U2585 partially releases translation termination arrest on *tnaC*, in agreement with the proposed functional role for this residue^36^. Moreover, the efficiency of translation termination is known to be dependent on the sequence context around the C-terminal amino acid, with the penultimate amino acid playing a critical role during peptidyl-tRNA hydrolysis^37^. The TnaC-dependent inhibition of RF2 action may thus represent an extreme form of these context-dependent effects, in which TnaC residues 23 and 24 prevent the GGQ loop of RF2 from adopting a catalytically active conformation within the PTC, ultimately resulting in impaired translation termination and ribosome stalling.

## DISCUSSION

In this work we identified and characterized a TnaC variant with a single R23F mutation that undergoes translational arrest at much lower L-Trp concentrations than the wild-type TnaC. Our structural data reveal how the ribosome and nascent TnaC cooperate to capture a single molecule of L-Trp and explain the role of various functionally important residues, such as the strictly conserved Trp12 and Asp16 of TnaC, the 23S rRNA base pair A752:U2609 and residue Lys90 of uL22. Visualization of a stalled TnaC(R23F)–70S–RF2 complex yields fresh insights into the mechanism by which TnaC inhibits its own hydrolysis from tRNA^Pro^, with 23S rRNA residue U2585 and residues 23 and 24 of TnaC proposed to interfere with the correct folding of the RF2 catalytic loop. Unexpectedly, the wild-type and R23F TnaC–ribosome complexes exhibit nearly identical conformations of the nascent peptide and the ligand-binding site, indicating that an R23F mutation within a region of TnaC that is positioned near the PTC does not induce an allosteric conformational change resulting in a binding site with increased affinity for L-Trp. Instead, our data suggest that the competition between L-Trp binding and peptidyl tRNA hydrolysis (or puromycin cleavage) determines the tryptophan sensitivity of the system, and leads us to propose a model to explain tryptophan-dependent ribosome stalling on the last codon of *tnaC* (Fig. 6a).

**Figure 6.**
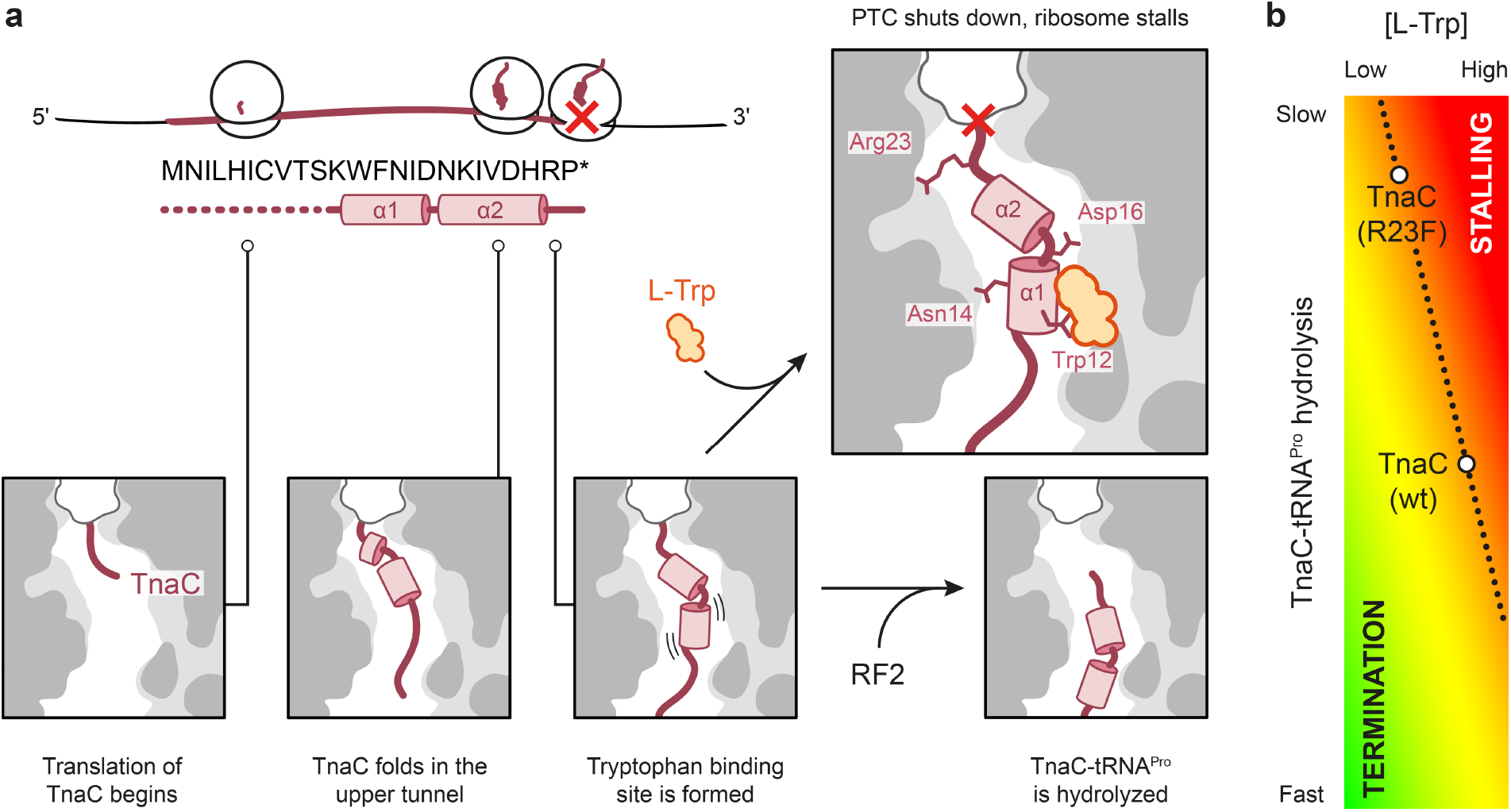
Mechanism of L-Trp sensing by a TnaC–ribosome complex. **a**, Model for the binding of L-Trp (orange) by the ribosome and TnaC (red), leading to inactivation of the peptide release activity of the ribosome (red cross). **b**, Schematic depicting the proposed relationship between the rate of TnaC-tRNA^Pro^ hydrolysis and the concentration of L-Trp required to achieve a given outcome, ranging from termination (green) to stalling (red) .

To begin, translation of the first ten amino acids of TnaC proceeds normally, with the nascent peptide progressing through the ribosomal exit tunnel unimpeded. Synthesis and folding of the α1 helical segment containing the WFNID motif ensues, followed by α2, eventually resulting in a fully synthesized peptide with a compact C-terminus similar to that observed in our tryptophan-containing complexes (Fig. 6a). This structure would make it difficult for the C-terminal region of nascent TnaC to pass through the tunnel constriction and would partially interfere with the correct positioning of the GGQ loop of RF2, giving rise to the tryptophan-independent pause thought to be responsible for the basal response *in vivo*^14^. Since the R23F mutation resulting in increased tryptophan sensitivity is located at the time of termination arrest near the PTC and does not alter the conformation of the L-Trp binding site, we propose that the kinetics of peptide release could be the basis of this phenotype (Fig. 6b). This hypothesis is supported by the previous observation that RF2 protein variants that are known to be less effective at catalyzing hydrolysis are better inhibited by L-Trp^33^. For this to be the case, RF2 would have to compete with L-Trp for binding to the ribosome, implying that ligand binding would only occur once TnaC synthesis is complete. Such a late binding event is consistent with our structural data. First, the WFNID motif of TnaC must fold into helical segment α1 prior to binding the L-Trp ligand. Second, α1 must not cross the tunnel constriction if it is to engage with an L-Trp molecule located next to the U2609:A752 base pair. Third, α-helices are known to fold on the nanosecond timescale in solution^38^, some three orders of magnitude faster than the incorporation of an amino acid into the nascent chain, meaning that α2 would also be folded by the time α1 reaches the constriction and binds L-Trp. Fourth and last, the wide channel leading to the ligand binding site in our structures is compatible with the late arrival of L-Trp. Thus, it appears that the system would only be primed for ligand binding once the ribosomal P-site is positioned on codon 23 or 24 of *tnaC* and the nascent peptide is fully synthesized.

Because the R23F mutant requires a much lower concentration of L-Trp to achieve the same response as wild-type TnaC, our model implies that the susceptibility of the tryptophan-free TnaC(R23F)–70S complex to RF2 action must be lower than that of the wild-type TnaC–70S complex (Fig. 1f and Fig. 6b). Such differences in RF2 susceptibility in the absence of bound L-Trp could be explained structurally by the extent to which the Arg and Phe amino acid side chains at position 23 of TnaC prevent the GGQ loop of RF2 from reaching its active conformation. A more severe steric block caused by the Phe23 side chain of the R23F variant may result in an increased dwell time of the tryptophan-free TnaC(R23F)–70S complex on codon 24 of *tnaC*, giving additional time for L-Trp to bind and negating the need for a high concentration of ligand to achieve the same response as wild-type TnaC.

Only a handful of other ligand-dependent arrest peptides have been structurally characterized to date^26,39–42^, including only one other bacterial amino acid-sensing peptide^26^. The best-characterized group comprises drug-dependent arrest peptides belonging to the Erm family, which undergo translational arrest in response to macrolide antibiotics to regulate the expression of macrolide-resistance genes. These arrest peptides, among which ErmBL^42^ and ErmCL^41^ have been characterized structurally, constitute a special case in that the antibiotic ligand binds to the ribosome with high affinity even in the absence of nascent peptide^30^. As a result, macrolide-dependent arrest peptides are akin to ligand-independent arrest peptides like SecM^43^, MifM^44^ or VemP^27^, with the difference that the shape and dimensions of the exit tunnel are altered by the bound ligand. In contrast, the low affinity of L-ornithine for the ribosome requires that SpeFL, an arrest peptide found in certain types of γ-proteobacteria, use a distinct mechanism to sense this non-proteinogenic amino acid^26^. In this case, the N-terminal sensor domain of SpeFL folds first, enabling ligand recognition by the ribosome and the nascent peptide after it has crossed the tunnel constriction. Synthesis and compaction of the C-terminal effector domain ensues, leading to the inhibition of RF1-mediated peptide release. Although both wild-type TnaC and SpeFL undergo termination arrest and require high ligand concentrations to ensure that binding prevails over continuing translation, the mechanism by which they do so differs. SpeFL binds L-ornithine long before peptide synthesis is complete, whereas TnaC must be fully formed for L-Trp binding to occur. In the former case, L-ornithine is thought to pre-associate with the ribosomal exit tunnel, whereas L-Trp binding appears to be a late event that determines the fate between peptide release and ribosome stalling. In both cases however, the peptide and the ribosome co-operate to form the ligand binding site and the dynamics of the system are paramount. It is likely that variants of the mechanisms employed by TnaC or SpeFL are also used by other arrest peptides to sense small molecular weight ligands with low affinity for the ribosome^45^, such as the arginine attenuator peptide in *Neurospora crassa*^46^ or the sucrose-sensing uORF2 peptide from *Arabidopsis thaliana*^47^. Finally, the existence of a tryptophan-independent pause during the translation of *tnaC* immediately prior to ligand binding suggests that certain ligand-dependent arrest peptides may have evolved from constitutive arrest sequences. It is therefore possible that a fraction of the many small ORFs populating bacterial and eukaryotic genomes^48^ encode peptides capable of transiently arresting the ribosome, and that these small ORFs may act as reservoirs from which metabolite-dependent arrest peptides could arise to adapt to new selective pressures.

## Acknowledgements

A.-X.S., A.H.V. and C.A.I. have received funding for this project from the European Research Council (ERC) under the European Union’s Horizon 2020 research and innovation program (Grant Agreement No. 724040). C.A.I. is an EMBO YIP and has received funding from the Fondation Bettencourt-Schueller. We acknowledge the European Synchrotron Radiation Facility for provision of microscope time on CM01^62^ and we thank Eaazhisai Kandiah for her assistance. We thank Yaser Hashem for help with grid preparation and Pierre Nottelet for help with data collection on the Talos Arctica microscope at the European Institute for Chemistry and Biology. L.R.C.-V. and N.V.-L. received funding for this work from the National Science Foundation U. S. A. [MCB-1158271 to L.R.C.-V. and MCB-1951405 to N.V-L.] and M.S.S. from the National Institutes of Health [R01 GM47498].

## Author contributions

L.R.C.-V., C.A.I., M.S.S. and N.V.-L. designed the study. A.K.M. isolated the loss-of function suppressors. A.S and D.K. generated toe-printing analysis. L.R.C.-V. performed the Western blots and Northern blots used to detect components of isolated arrested ribosomes. E.R.G performed the *in vitro* TnaC-tRNA accumulation analysis and puromycin cleavage assays. A.H.V. designed the wild-type TnaC construct and carried out initial complex purification. A.-X.S. designed the R23F construct and performed complex preparation. T.N.P. prepared the cryo-EM grids and performed data collection on the Talos Arctica. A.-X.S. processed the cryo-EM data and built the models. A.-X.S., C.A.I. and L.R.C.-V. wrote the paper. A.-X.S., C.A.I., L.R.C.-V., M.S.S. and N.V.-L. reviewed and edited the manuscript.

## Competing financial interests

The authors declare no competing financial interests. Correspondence and requests for materials should be addressed to C.A.I., L.R.C. or M.S.S.

## METHODS

### Toeprinting assays

Toeprinting assays shown in Fig. 1b and Fig. 1d were performed as previously described^25^. Customized PURExpress reconstituted systems lacking all amino acids and release factors (Δaa, ΔRF123) (New England Biolabs) (Fig. 1b) or a complete system (New England Biolabs) (Fig. 1d) were used to perform transcription-translation reactions. DNA fragments containing wild-type and mutant alleles of the *tnaC* gene were obtained by polymerase chain reaction (PCR) using the plasmids and oligonucleotides indicated in Supplementary Table 3. Coupled transcription-translation reactions (5 μl) were carried out with 0.1 to 0.3 pmol/μl *tnaC* DNA fragments. Variable concentrations of L-Trp (0, 0.02, 0.04, 0.08, 0.16, 0.32, 0.64, 1.25, 2.5, 5 and 10 mM) were used for the toeprint in Fig. 1b, whereas 0.3 mM or 4 mM L-Trp was used for the experiments shown in Fig. 1d. The 19 other amino acids were supplemented to a final concentration of 0.3 mM in all cases. Reaction mixtures were incubated at 37°C for 30 minutes. cDNA was synthesized by adding 4 units of AMV reverse transcriptase enzyme (Roche) and 0.5 μl 20 pmol/μl of reverse primer labeled with [^32^P] per reaction. Reactions were then incubated at 37°C for 15 min. cDNA products were resolved by electrophoresis using 6% urea-polyacrylamide gels. Final gels were exposed to a storage phosphor screen for 1 hour and scanned using a Typhoon Imager 9410 (GE Healthcare). Band intensities were determined using the Fiji ImageJ software^49^. To produce the bar plot shown in Fig. 1c, densitometry values obtained with the Fiji ImageJ software were plotted to calculate the percent accumulation of arrested ribosomes with the following formula:

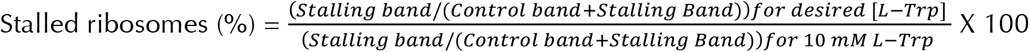

where a constant signal band located 20 nucleotides downstream of the stalling bands was used as a loading control band. Calculated values were plotted in GraphPad Prism 6 with one site specific binding (hyperbola) curve fitting 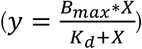 with both B_max_ and K_d_constrained to be greater than zero.

### TnaC-tRNA^Pro^ cleavage assays

To determine the accessibility of TnaC-tRNA^Pro^ for puromycin induced cleavage (Fig. 1e), *in vitro* isolated complexes were obtained using wild-type and mutant *tnaC* biotinylated-mRNAs as indicated previously^16^. In short, 2 µg of wild-type or *tnaC* mutant biotinylated-mRNAs obtained from PCR fragments produced with the plasmids and oligonucleotides indicated in Supplementary Table 3 were added to 25 µl *in vitro* reactions consisting of a ΔRF123 cell-free extract supplemented with [S^35^]-L-methionine and 4 mM L-Trp. After incubation at 37°C for 10 min, complexes were isolated using streptavidin-paramagnetic beads, and washed with buffer containing 35 mM Tris-HCl pH 8.0, 10 mM magnesium acetate, 175 mM potassium glutamate, 10 mM ammonium acetate and 1 mM dithiothreitol. Isolated complexes devoid of L-Trp were then incubated at 37°C following the sequential addition of 4 mM L-Trp for 2 min, and 1 mM puromycin for an additional 10 min. Reactions were stopped, resolved in 10% Tris-tricine polyacrylamide gels, and [S^35^]-labeled products were quantified as described above using the Fiji ImageJ software. To obtain the bar plot shown in Fig. 1f, the % of remaining TnaC-tRNA^Pro^ was calculated using the following formula:

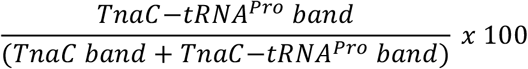

### Cryo-EM sample preparation

Bicistronic templates encoding two successive copies of wild-type *tnaC* or the *tnaC(R23F)* variant were purchased from Eurofins, France (Supplementary Table 3). Stalled TnaC–ribosome complexes were prepared as described previously^26^. Briefly, a 750 µL *in vitro* translation reaction (RTS100 HY kit; Biotechrabbit) was performed for 1 h at 30°C in the presence of 2 mM L-Trp, puromycin was added to 100 µM and the reaction was further incubated for 3 min. The sample was separated on 10-40% sucrose gradients in Buffer A (50 mM HEPES-KOH, pH 7.5, 100 mM potassium acetate, 25 mM magnesium acetate and 2 mM L-Trp) for 2h 45min at 35,000 rpm in a TH641 rotor (Thermo) and polysomal fractions were collected using an ultraviolet detection system (UA-6; Teledyne ISCO) coupled to a gradient fractionator (Foxy R1; Teledyne ISCO). Sucrose was removed by 7 sequential washes in a centrifugal filtration unit with a molecular weight cut-off (MWCO) of 100 kDa (Millipore). Polysomes were converted into monosomes by RNase H digestion, separating the two consecutive copies of *tnaC* on the mRNA. This was achieved by mixing 50 µL of polysomes with 50 µL of a 100 µM stock of RNase H oligonucleotide (Supplementary Table 3), 4 µL RNase H (New England Biolabs), 644 µL buffer A and 2 µL DTT (1M). The reaction was incubated for 1 h at 25°C. Sucrose gradients and removal of the sucrose were performed as described above, to collect the monosome fraction enriched in stalled TnaC–ribosome complexes.

### Cryo-EM grid preparation and data acquisition

Stalled TnaC–ribosome complexes were diluted to 200 nM in buffer A and 5 µL of this dilution were applied to Quantifoil (QF-R2/2-Cu) carbon grids, after these were coated with a 2 µm carbon layer using an CCU-010 Compact Coating Unit (Safematic) and glow discharged for 20 s at 2 mA. Grids were plunge-frozen in liquid ethane using a Vitrobot (FEI) set to 4°C and 100% humidity with 2.5 s blotting time and 30 s wait time. TnaC and TnaC(R23F)_B grids were imaged using a 200 kV Talos Arctica microscope (FEI), equipped with a K2 Summit direct electron detector (Gatan) in nanoprobe mode. Images were recorded with SerialEM^50^ in counting mode with a magnified pixel size of 0.93 Å at a magnification of 45,000x. The defocus range was set to -1.0 to -2.0 or -0.4 to -2.0 µm for the TnaC (IECB) and TnaC(R23F)_B (IECB) data sets, respectively, and each movie contained 38 frames with a dose per frame of 1.31 e/Å^2^. TnaC(R23F)_A grids were imaged on a 300 kV Titan Krios microscope (FEI), equipped with a K2 Summit direct electron detector (Gatan). Images were recorded with SerialEM^50^ in counting mode with a magnified pixel size of 0.827 Å at a magnification of 165,000x. The defocus range -0.4 to -1.6 µm and each movie contained 40 frames with a dose per frame of 1.1 e/Å^2^.

### Cryo-EM data processing

Raw micrographs were processed using *RELION v*.*3*.*1*^51^ as depicted in Supplementary Fig. 2. Briefly, movie frames were aligned using RELION’s implementation of *MotionCor2*^52^, and *Gctf*^53^ was used for Contrast Transfer Function (CTF) estimation. After filtering out bad micrographs by resolution, 2 rounds of 2D classification were performed on 4x downscaled particles in *RELION v*.*1*.*3*^51^. Unsupervised 3D classification was performed to eliminate the ratcheted ribosome class, as well as junk particles. Focused 3D classification on the A-, P- and E-tRNA sites was carried out with 4x downscaled particles to select classes with an occupied P-site or P/E sites. These particles were pooled, re-extracted at full size (408px or 464px) and polished using iterative Bayesian polishing and CTF refinement on a per particle basis. These ‘shiny’ particles were refined, leading to the final 2.9 Å and 2.4 Å maps of the TnaC–70S and TnaC(R23F)–70S complexes, respectively. During the last refinement step, a mask covering the large subunit was used at each cycle to align all particles against this subunit. In the case of the TnaC(R23F)–70S–RF2 complex, particles from two datasets (TnaC(R23F)_A and TnaC(R23F)_B) collected on different microscopes were merged according to the procedure described in ref.^54^. First, both datasets were refined independently, and pixel sizes were calibrated on the basis of the correlation between a simulated map obtained from a preliminary model and the refined cryo-EM density map. Polished particles from the TnaC(R23F)_A (ESRF) dataset (Titan Krios) were then resized to match those of the TnaC(R23F)_B (IECB) particles (Talos Arctica), CTF values were adjusted and the rescaled particles were merged with the polished TnaC(R23F)_B (IECB) particles, yielding a final reconstruction with an overall resolution of 2.6 Å resolution.

### Model building and refinement

Density maps were sharpened with *Phenix Autosharpen*^55^ without a model, and the pixel size was optimized using the *ChimeraX*^56^ fit-in-map feature by comparison to a simulated map calculated from PDB 6TBV^26^. The same atomic coordinates were also used for the initial ribosome model, while tRNA^Pro^, the TnaC nascent peptide and the L-Trp ligand were manually built in *Coot*^57^ and improved through iterative rebuilding/refinement in *Phenix Real Space Refine*^58^ and *ISOLDE*^59^. In the case of the TnaC(R23F)–70S–RF2 structure, the initial molecular model of RF2 was obtained from PDB 6OUO^60^. Water molecules were added to the TnaC(R23F)–70S model using the *douse* algorithm of *Phenix*^61^.

### Figure preparation

Depictions of the map densities and molecular models were made using *ChimeraX*^56^ and *Pymol* open-source (Schrödinger, LCC).

### Western and Northern blots

For Fig. 5c, isolated arrested ribosomes were obtained as indicated above using a ΔRF123 cell-free extract. Reactions performed in the presence of 20 nM of each RF2 variant were resolved by electrophoresis on 10% Tris-tricine gels and transferred to nylon membranes. RF2 protein variants were produced from bacteria expressing the plasmids indicated in Supplementary Table 3 and purified as described previously^33^. rRNA from the isolated complexes were obtained using standard phenol-chloroform extractions and resolved in 1% non-denaturing agarose gels.

### Data availability

The TnaC–70S, TnaC(R23F)–70S and TnaC(R23F)–70S–RF2 structures have been deposited with the Research Collaboratory for Structural Bioinformatics Protein Data Bank with accession codes 7O19, 7O1A and 7O1C; the cryo-EM maps have been deposited with the Electron Microscopy Data Bank with accession codes EMD-12693, EMD-12694 and EMD-12695. Raw movie stacks have been deposited with the Electron Microscopy Public Image Archive with accession codes EMPIAR-XXXXX, EMPIAR-XXXXX and EMPIAR-XXXXX. Source data for Fig. 1, Fig. 5, Supplementary Fig. 1 and Supplementary Fig. 4 are provided with the paper.

## SUPPLEMENTARY INFORMATION

### SUPPLEMENTARY NOTES

#### Isolation of suppressors of the loss-of-function mutant *tnaC* (D16E) gene

Intragenic suppressor mutations to the loss-of-function *tnaC* D16E gene were obtained using the strain AW845 (MG1655 Δ7 *rrn* Δ(*lacZYA*) Δ(*tnaCAB*) *att7*::*tna*_*p*_*tnaC*(*D16E*)(*tnaA*’-‘*lacZYA*) (p*rrnC*-*sacB*, ptRNA67), which carries a *tnaA*–*lacZ* protein fusion whose expression is under the control of the *tna* operon regulatory leader region (Ref. 15). This bacterial strain was grown overnight in Vogel Bonner (VB) minimal medium supplemented with 0.2% glucose, 0.05% acid-hydrolyzed casein, 0.01% vitamin B1 and 25 μg/ml kanamycin. Cells were washed once in VB without any supplements and 100 μl of washed cells were plated on VB minimal medium supplemented with 1% lactose, 100 μg/ml L-Trp, 0.01% vitamin B1 and 25 μg/ml kanamycin. Plates were incubated at 37°C for 6 days. Colonies that formed were picked and each colony was grown overnight in VB minimal medium supplemented with 0.2% glucose, 0.05% acid-hydrolyzed casein, 0.01% vitamin B1 and 25 μg/ml kanamycin. Since mutations that restore induction to the uninducible TnaC(D16E) mutant were desired, these cultures were screened for L-Trp inducibility. Overnight cultures were replica plated onto VB plates supplemented with 1% lactose, 0.01% vitamin B1, and 25 μg/ml kanamycin with or without 100 μg/ml Trp. Colonies that either grew on the L-Trp containing plate and not at all on the plate lacking L-Trp, or those that grew better on the L-Trp containing plate than they did on the plate lacking L-Trp were selected for further analysis by standard Miller Assay. Out of the 1,152 total colonies screened by replica plating, 111 showing differential growth were analyzed by Miller Assay of exponentially growing cultures to determine if the LacZ reporter was inducible by L-Trp. Out of these 111 colonies, 13 were confirmed to be inducible by L-Trp in three independent experiments. The leader region of the reporter operon from the 13 mutants was sequenced to determine if there were any mutations within *tnaC* using primers listed in Supplementary Table 3. Based on the *tnaC* sequence, the mutants were divided into 4 classes. Nearly half of the mutants were true-revertants, which means that the combined selection and screening method is a powerful tool that can be used to identify mutations that restore L-Trp-inducibility to the non-functional D16E mutant, or in principle any other non-functional TnaC mutant. Mutations at two different non-conserved codon positions within *tnaC* were found to suppress the non-functional D16E replacement. Mutants have either a second-site amino acid replacement S10P or at the residue R23 (R23S or R23H)). To ensure that the second-site mutations, S10P and R23H, within *tnaC* were responsible for restoring L-Trp-mediated ribosome arrest, strains containing reporter operons with the double changes S10P/D16E and D16E/R23H in *tnaC* were constructed on a MG1655 Δ(*lacZYA*) strain (Ref. 15) by using the plasmids and primers indicated in Supplementary Table 3. Derivates lacking an AUG start codon were also constructed to determine if the effects of these changes on the expression of the reporter genes were dependent or not on the translation of the *tnaC* gene variants. Expression levels of all these constructs are shown in Supplementary Table 1.

### Determination of the importance of the amino terminal end of TnaC on ribosome stalling

To determine the minimal elements of the TnaC peptide required for ribosome stalling we performed *in vitro* coupled transcription-translation assays (Ref. 15) using several *tnaC* constructs obtained by PCR that differ in the 5’-end of their open reading frame, generating N-terminally truncated versions of the TnaC peptide (Supplementary Fig. 4a). Accumulation of [^35^S]-TnaC-tRNA^Pro^, an indicator of ribosome stalling, was determined by resolving the reaction products in 10% Tris-tricine electrophoresis gels (Ref. 15). We observed that a deletion of the segment of amino acids 2-8 do not affect the accumulation of [^35^S]-TnaC-tRNA^Pro^ at higher L-Trp concentrations (5 mM), but is less sensitive to lower concentrations of L-Trp (Supplementary Fig. 4b, compare data for TnaC-16 with TnaC-24). However, deletion of residues 2–10 abolished the accumulation of [^35^S]-TnaC-tRNA^Pro^ (Supplementary Fig. 4b, compare data for TnaC-15 with TnaC-24). These results were corroborated using toeprinting assays (Ref. 15), which showed that deletion of residues 2–9 still produces a functional TnaC peptide at higher concentrations of L-Trp (Supplementary Fig. 4b, compare data for TnaC-15 with TnaC-24). None of the shorter peptides were functional (Supplementary Fig. 4b, compare data for TnaC-14 through TnaC-7 with TnaC-24). Overall, these results indicate that residues 10–24 contain enough structural information to detect L-Trp and inhibit translation termination.

**Supplementary Table 1.**
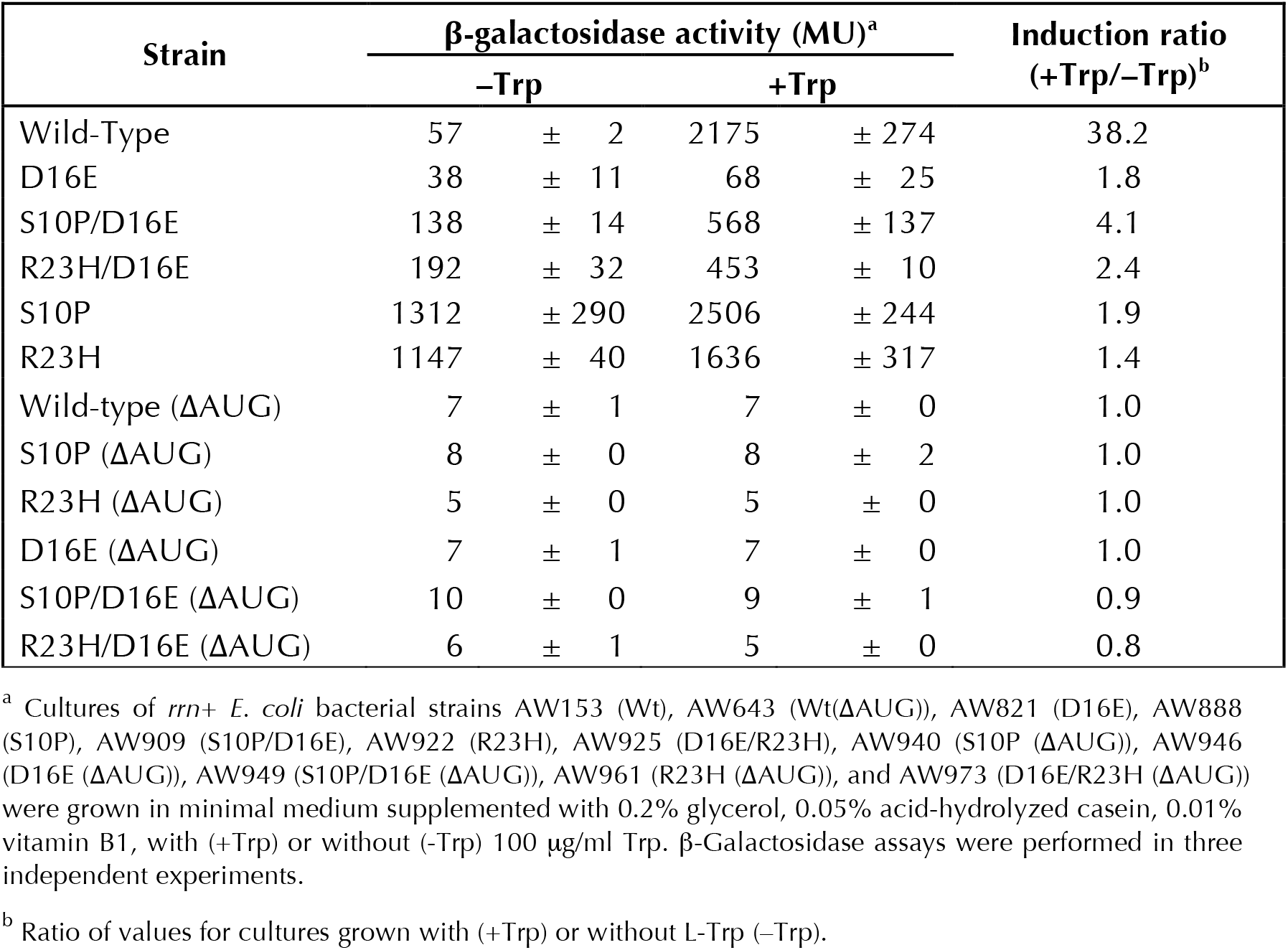
S10P and R23H are *cis*-acting mutations that suppress the loss-of-function D16E mutation.

**Supplementary Table 2.**
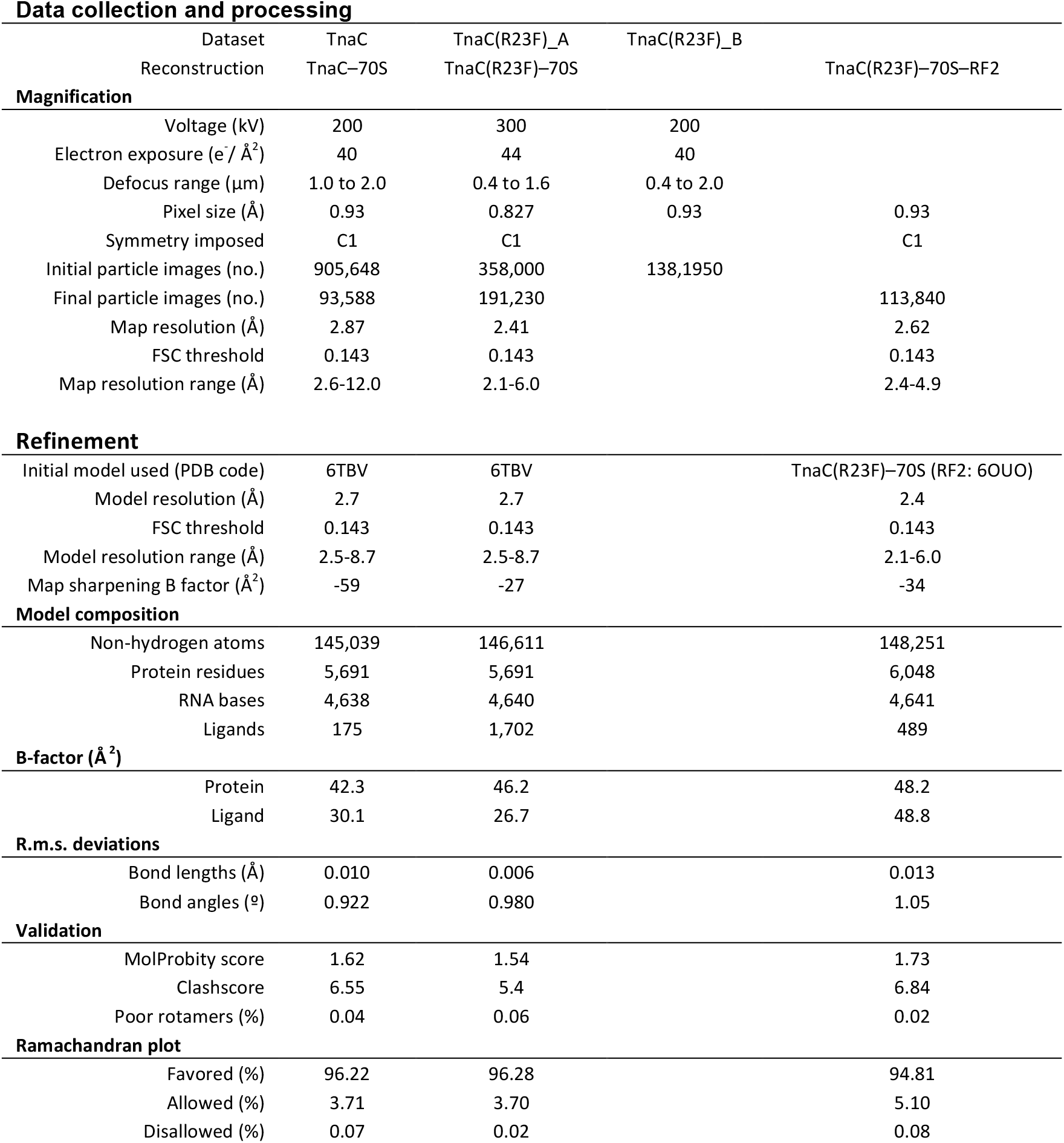
Cryo-EM statistics.

**Supplementary Table 3.**
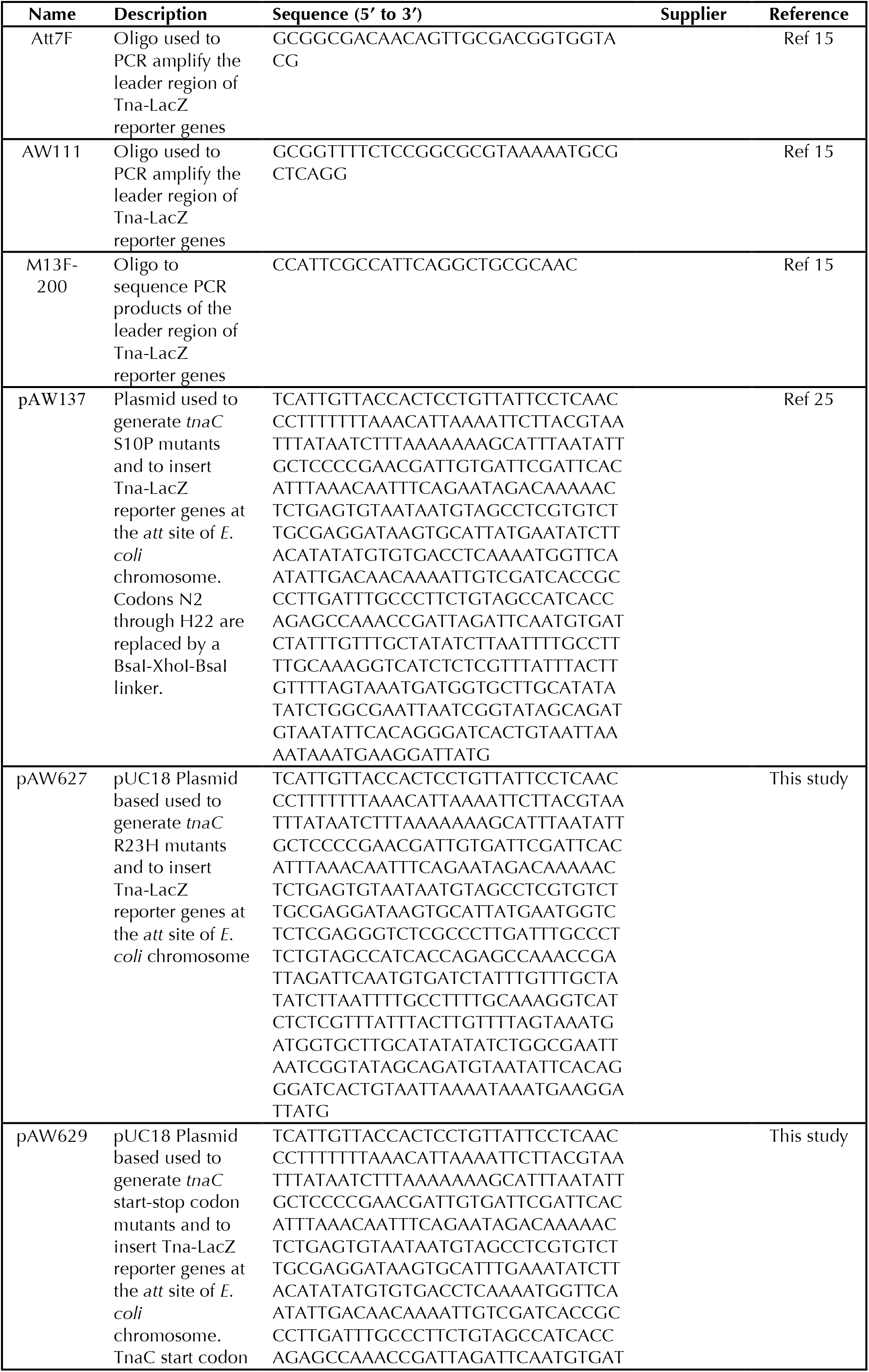
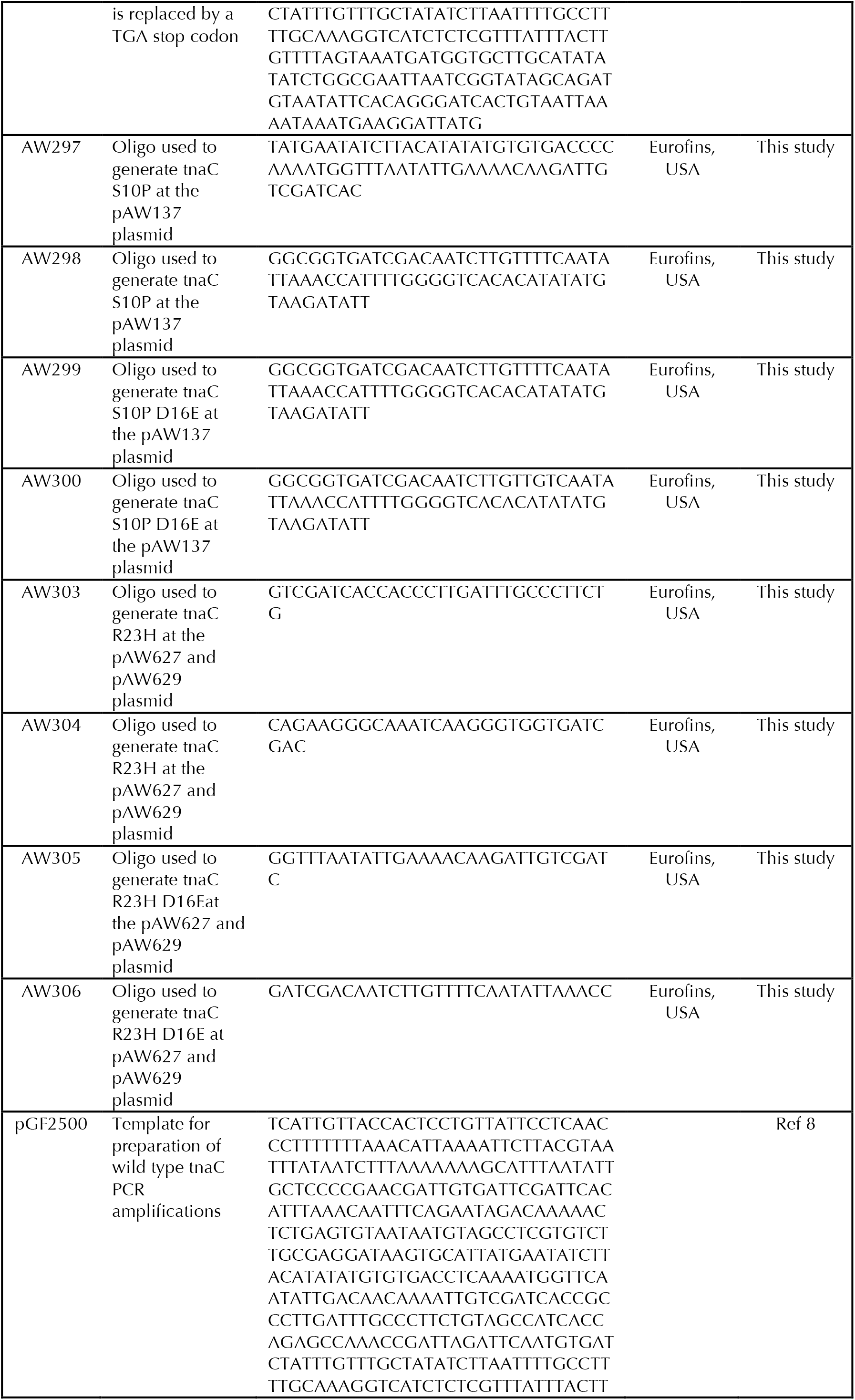
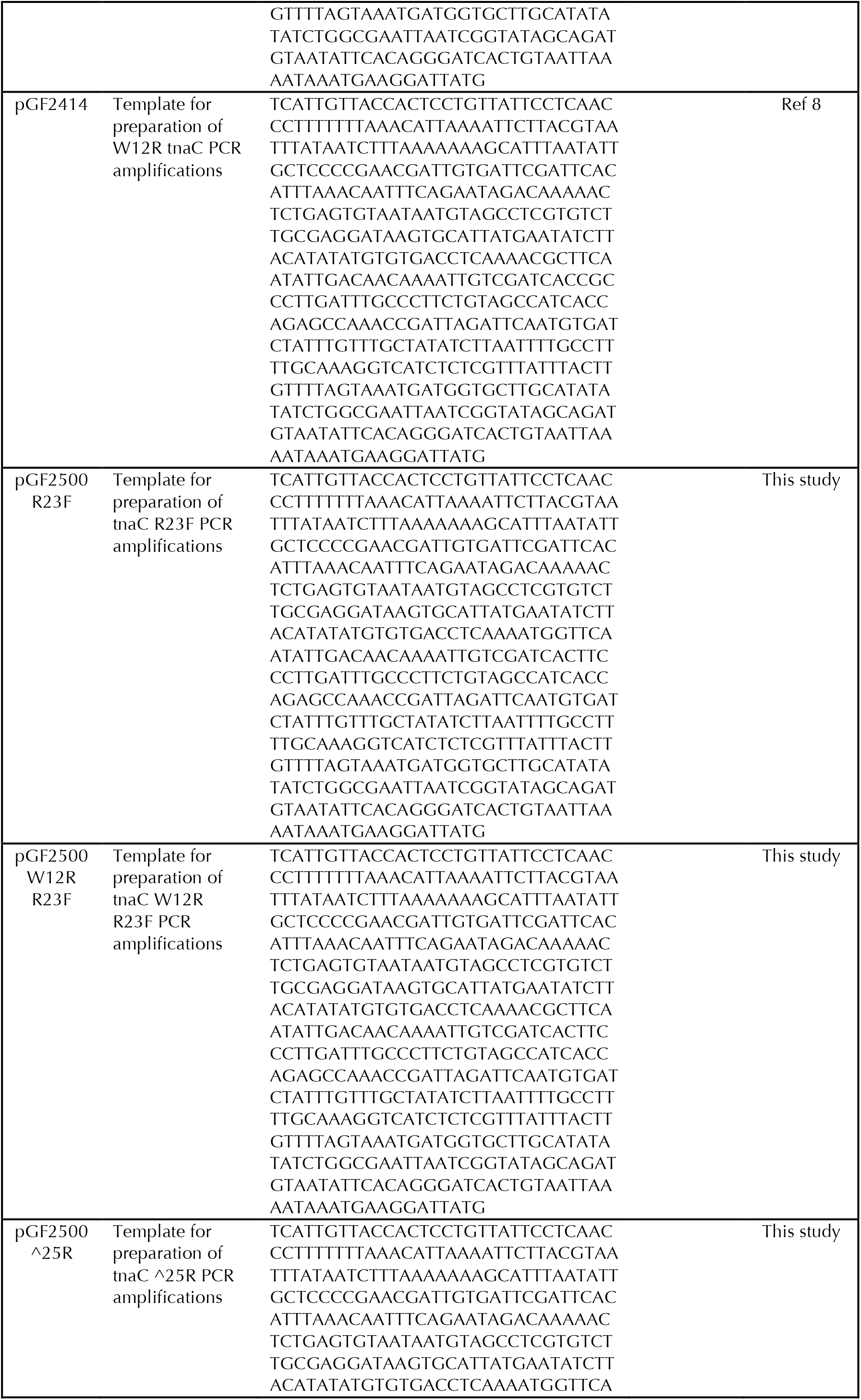
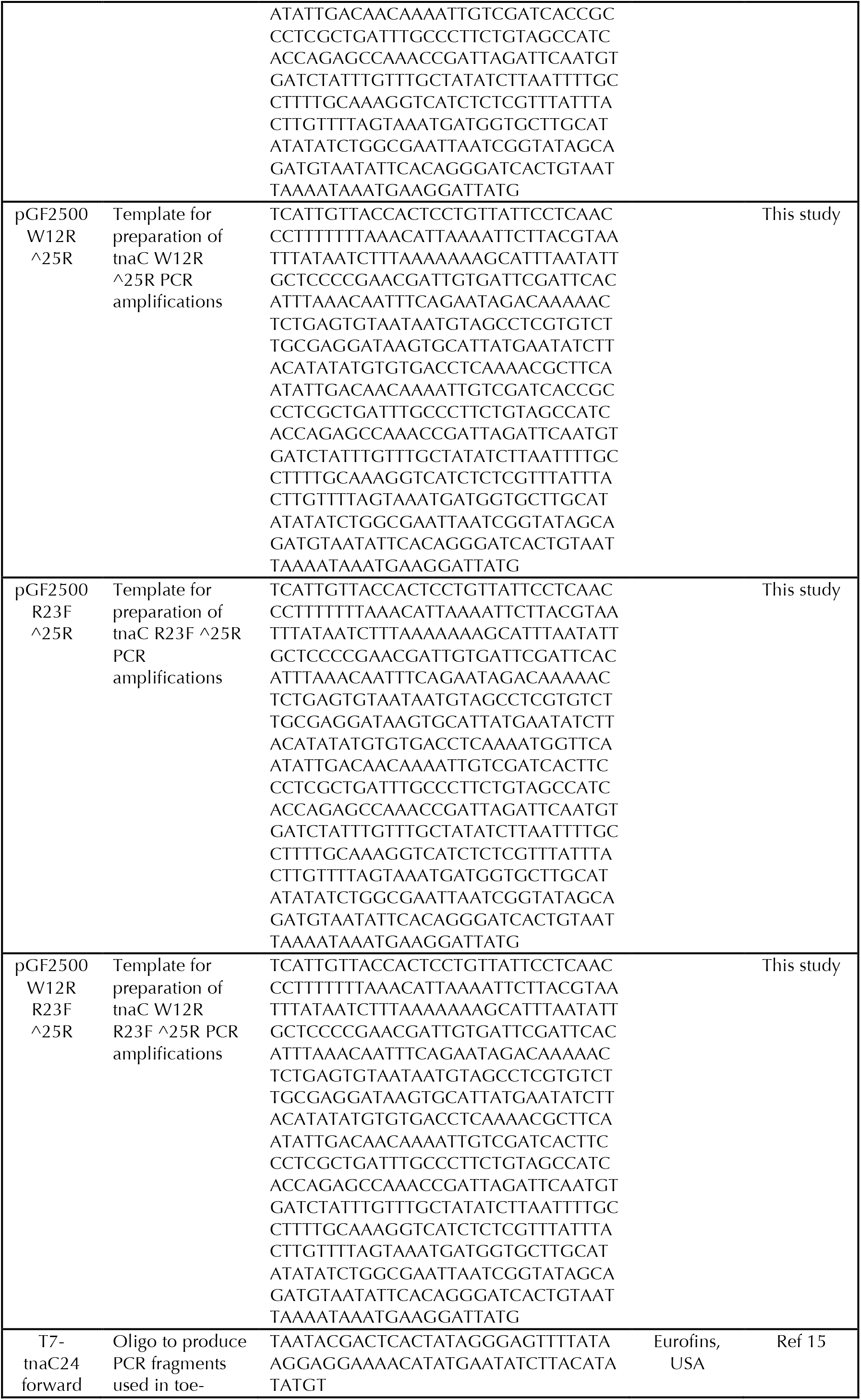
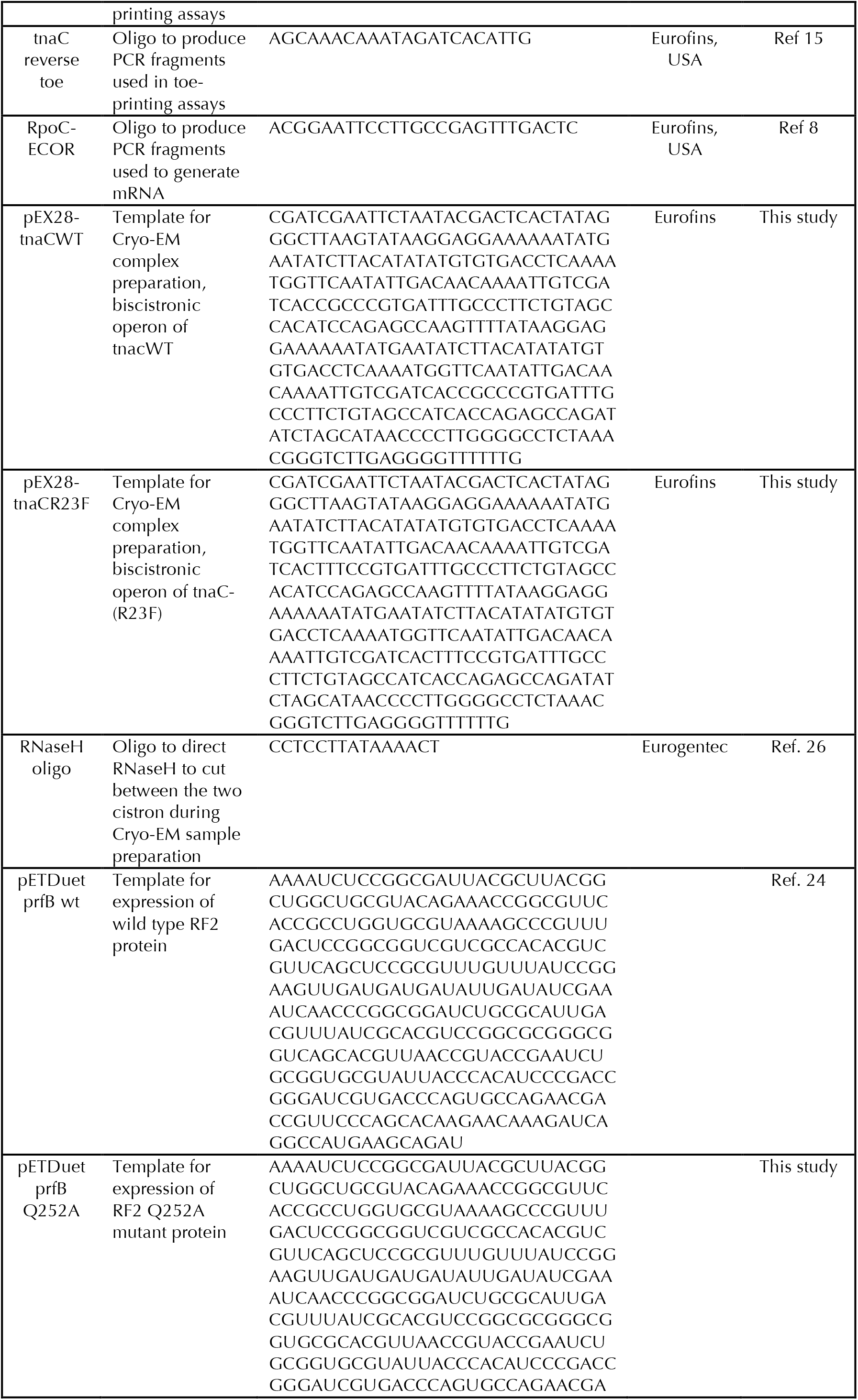
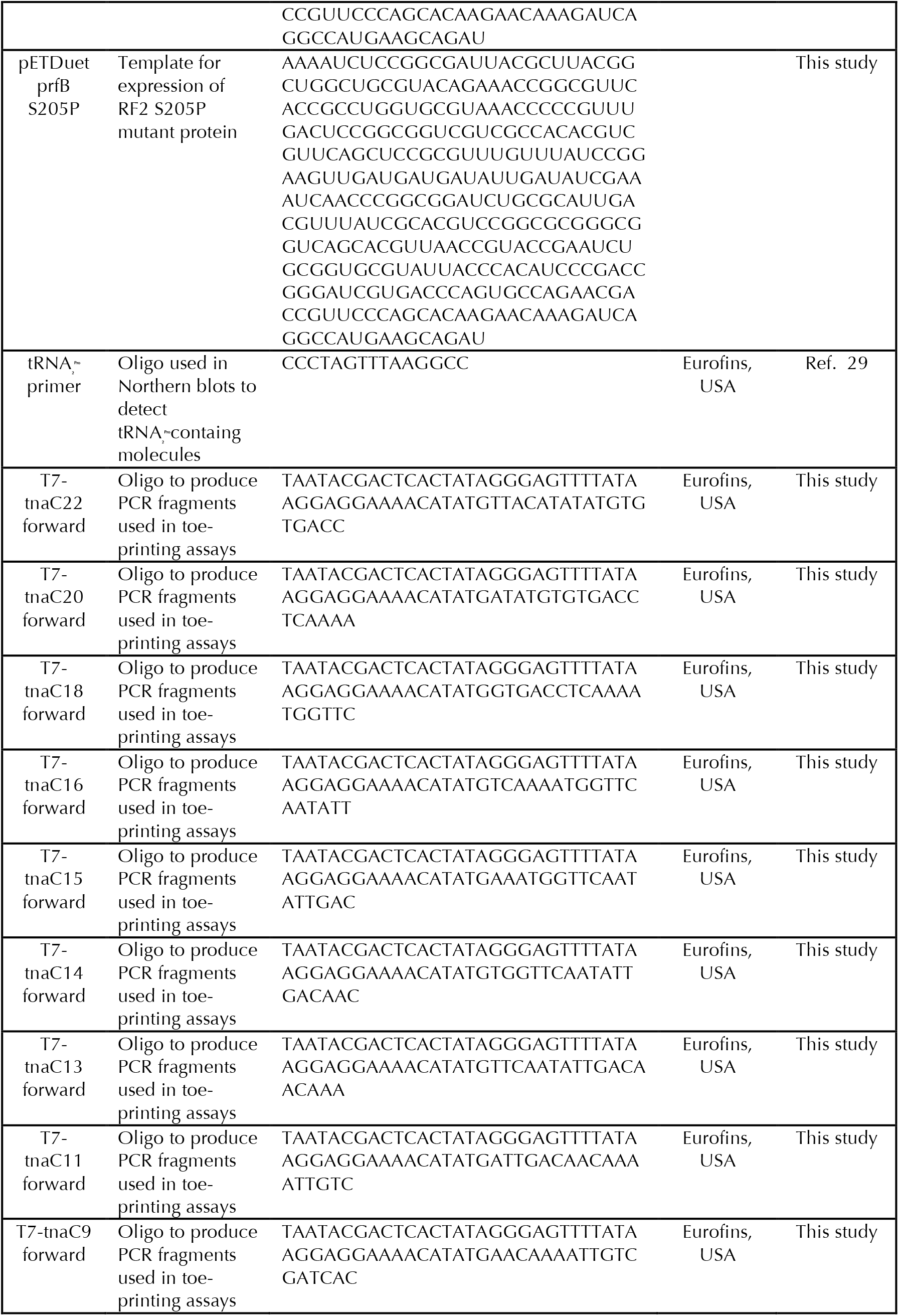
Constructs and oligonucleotides.

**Supplementary Figure 1.**
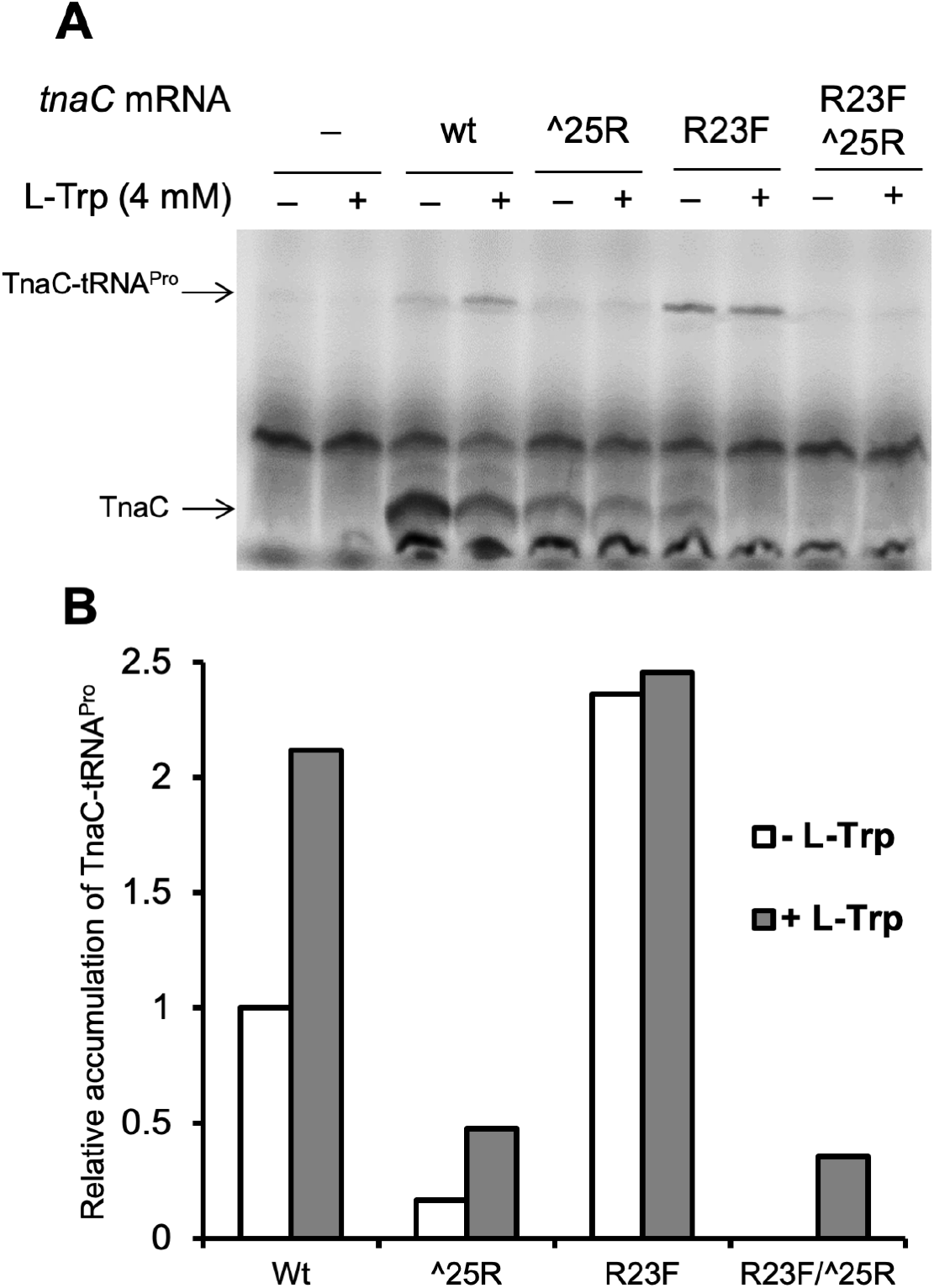
*In vitro* accumulation of TnaC-tRNA^Pro^ produced by several tnaC mRNA variants. (A) X-ray radiogram of the resolved products of *in vitro* translation assays performed in the presence of [^35^S]-labeled methionine, using mRNA templates encoding *tnaC* variants featuring arginine (wt) or phenylalanine (R23F) codons at position 23, an arginine codon at position 25 (^25R) or the double mutation R23F/^25R. The positions of [^35^S]-labeled free TnaC peptide and TnaC-tRNA^Pro^ are indicated. (B) Bar plot indicating the relative accumulation of TnaC-tRNA^Pro^ in the wild type and R23F mutant complexes. This experiment was repeated three times independently with similar results.

**Supplementary Figure 2.**
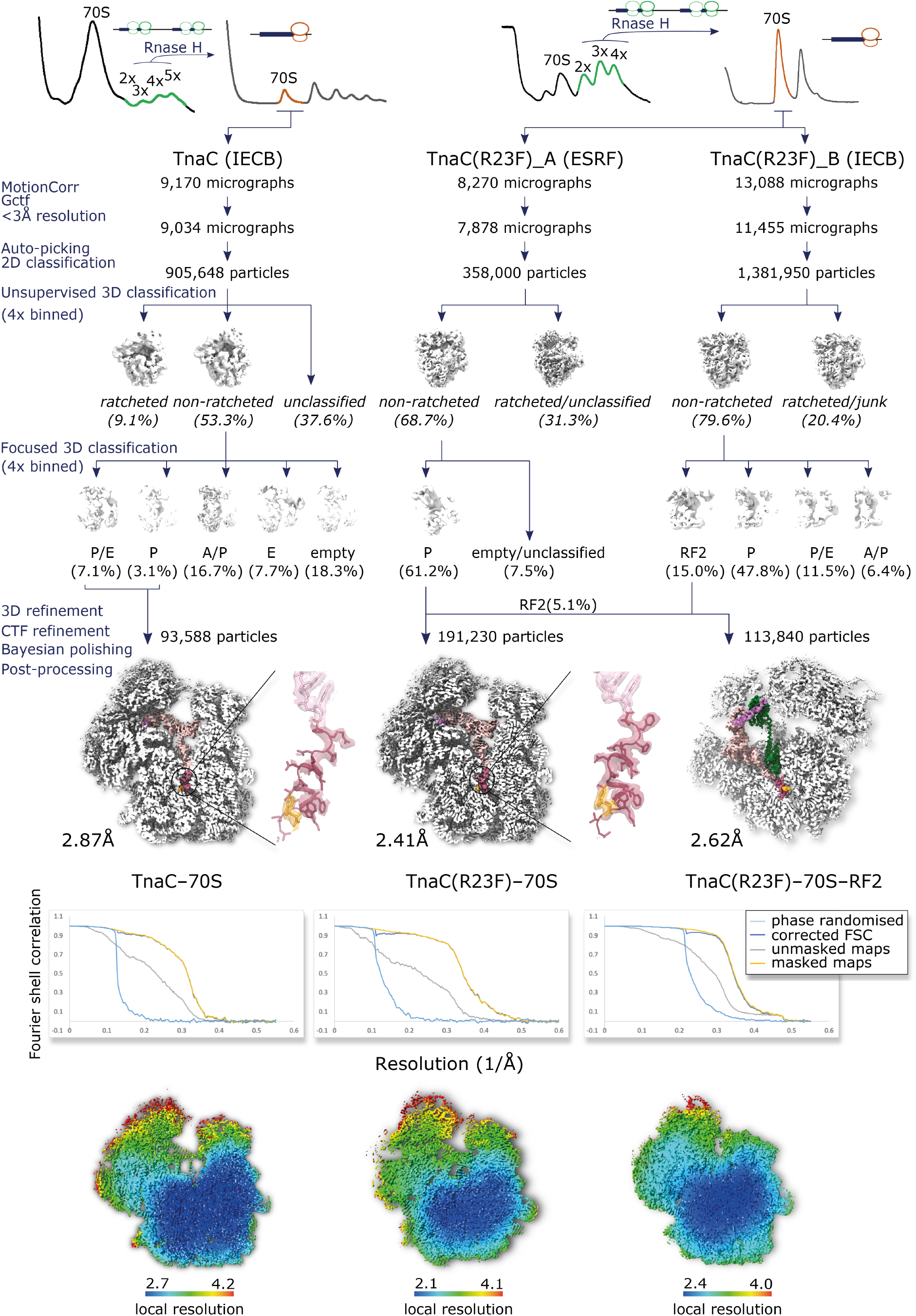
Complex purification and cryo-EM data processing workflow. Stalled ribosomal complexes were prepared using a bicistronic operon containing two identical copies of *tnaC* or *tnaC(R23F)*. A first sucrose gradient was performed to collect polysomes, followed by a second sucrose gradient after RNase H treatment to collect the monosomal fraction, which was the used to prepare the grids for cryo-EM data acquisition. The flowchart shows the workflow used to process and analyze cryo-EM data. Cross-sections of the final reconstructions are shown with the 70S ribosome in white, the mRNA in violet, the P-site tRNA in pink, the TnaC peptide in red and the L-Trp molecule in orange. Detailed maps of the TnaC peptide and L-Trp density are also shown for the TnaC–70S and TnaC(R23F)–70S complexes using the same color scheme, along with fitted atomic models. Fourier Shell Correlation (FSC) curves of the final reconstructions are shown as calculated by the *RELION 3*.*1* (Ref. 51) post-processing algorithm. Cross-sections of maps displaying the local resolution calculated by *RELION 3*.*1* (Ref. 51) are shown.

**Supplementary Figure 3.**
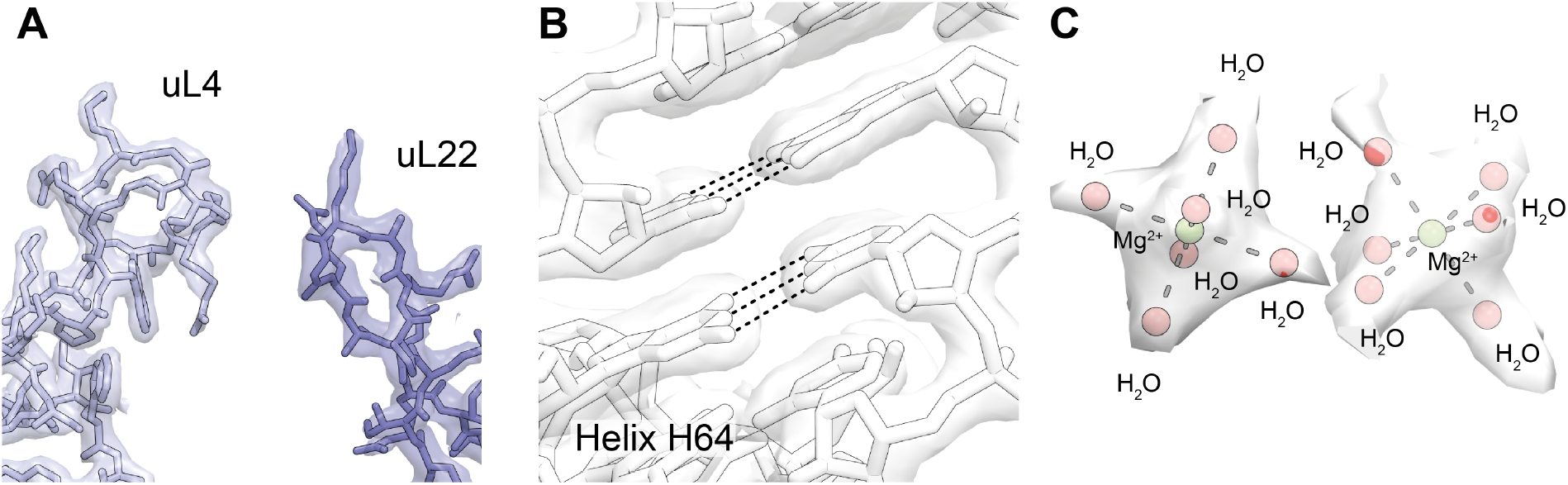
Quality of the cryo-EM density. Details of the TnaC(R23F)–70S map showing (A) the density and fitted atomic models for ribosomal proteins uL4 (light blue) and uL22 (periwinkle blue), near the ribosomal exit tunnel, (B) helix 64 of the 23S large subunit, and (C) two adjacent hydrated magnesium ion clusters in the core of the ribosomal large subunit.

**Supplementary Figure 4.**
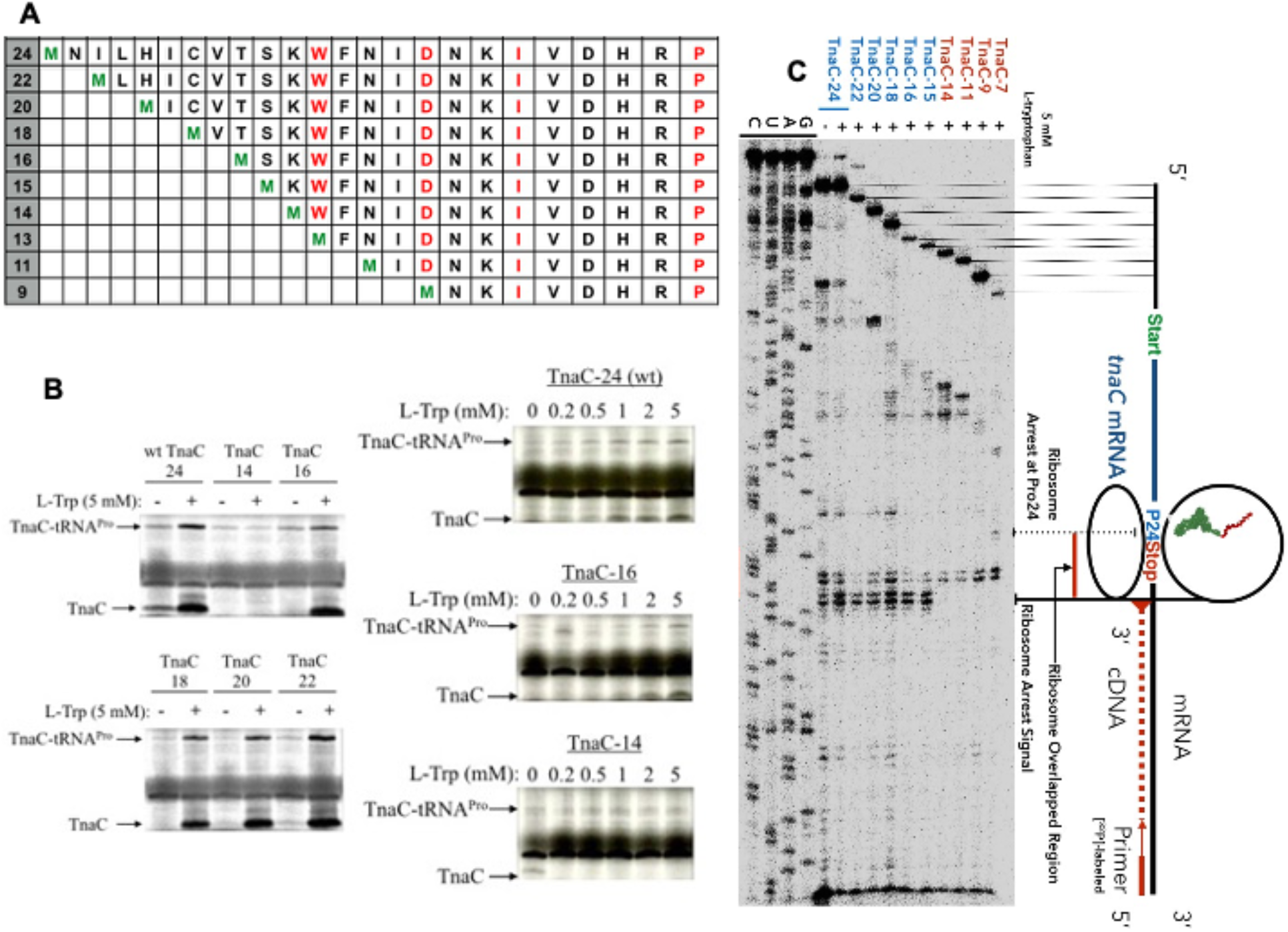
Effect of N-terminal deletions on the ability of TnaC to undergo translational arrest in the presence of L-Trp. (A) Diagram representing the sequences of the TnaC peptides used in the experiments shown in B and C. Green letters indicate the first decoded amino acid; red letters indicate conserved TnaC residues that have been shown experimentally to be functionally important. (B) Radiograms showing *in vitro* translation reactions performed as indicated before (Ref. 15). (C) Radiogram of a toe-printing assay performed as indicated before (Ref. 15). The schematic on the right shows the position on the *tnaC* mRNA of a ribosome stalled upon addition of L-Trp. The red dotted line indicates the cDNA product corresponding to the stalled ribosome in the toe-printing reaction.

**Supplementary Figure 5.**
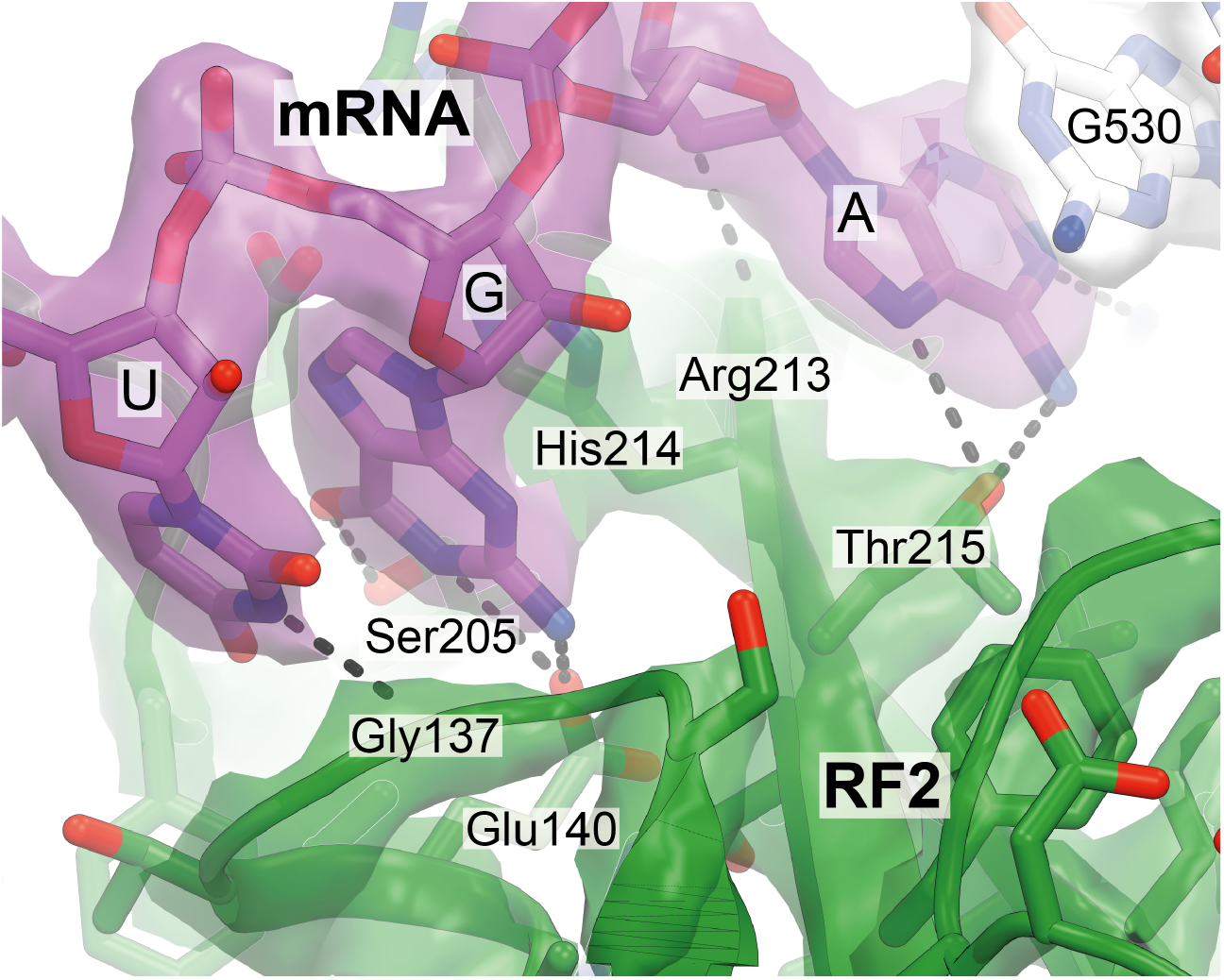
Interactions between RF2 and the UGA stop codon in the TnaC(R23F)–70S–RF2 structure. The stop codon in the A-site (purple) is recognized by domain II of RF2 (green).

## Notes

### Competing Interest Statement

The authors have declared no competing interest.

## REFERENCES

1. Edwards, R. M. & Yudkin, M. D. Location of the gene for the low-affinity tryptophan-specific permease of Escherichia coli. Biochem. J. 204, 617–619 (1982).

2. Deeley, M. C. & Yanofsky, C. Nucleotide Sequence of the Structural Gene for Tryptophanase of Escherichia coli K-12. J. Bacteriol. 147, 787–796 (1981).

3. Newton, W. A. & Snell, E. E. Catalytic Properties of Tryptophanase, a Multifunctional Pyridoxal. Proc. Natl. Acad. Sci. United States 51, 382–389 (1964).

4. Zarkan, A., Liu, J., Matuszewska, M., Gaimster, H. & Summers, D. K. Local and Universal Action: The Paradoxes of Indole Signalling in Bacteria. Trends Microbiol. 28, 566–577 (2020).

5. Botsford, J. L. & DeMoss, R. D. Catabolite repression of tryptophanase in Escherichia coli. J. Bacteriol. 105, 303–312 (1971).

6. Stewart, V. & Yanofsky, C. Evidence for transcription antitermination control of tryptophanase operon expression in Escherichia coli K-12. J. Bacteriol. 164, 731–740 (1985).

7. Konan, K. V. & Yanofsky, C. Rho-dependent transcription termination in the tna operon of Escherichia coli: Roles of the boxA sequence and the rut site. J. Bacteriol. 182, 3981– 3988 (2000).

8. Gong, F. & Yanofsky, C. Reproducing tna operon regulation in Vitro in an S-30 system: Tryptophan induction inhibits cleavage of TnaC peptidyl-tRNA. J. Biol. Chem. 276, 1974–1983 (2001).

9. Kamath, A. V. & Yanofsky, C. Roles of the tnaC-tnaA spacer region and rho factor in regulating expression of the tryptophanase operon of Proteus vulgaris. J. Bacteriol. 179, 1780–1786 (1997).

10. Gong, F. & Yanofsky, C. Instruction of translating ribosome by nascent peptide. Science 297, 1864–1867 (2002).

11. Gollnick, P. & Yanofsky, C. tRNA(Trp) translation of leader peptide codon 12 and other factors that regulate expression of the tryptophanase operon. J. Bacteriol. 172, 3100– 3107 (1990).

12. Gish, K. & Yanofsky, C. Evidence suggesting cis action by the TnaC leader peptide in regulating transcription attenuation in the tryptophanase operon of Escherichia coli. J. Bacteriol. 177, 7245–7254 (1995).

13. Konan, K. V. & Yanofsky, C. Regulation of the Escherichia coli tna operon: Nascent leader peptide control at the tnaC stop codon. J. Bacteriol. 179, 1774–1779 (1997).

14. Wang, T. et al. Dynamics of transcription–translation coordination tune bacterial indole signaling. Nat. Chem. Biol. 16, 440–449 (2020).

15. Martínez, A. K. et al. Crucial elements that maintain the interactions between the regulatory TnaC peptide and the ribosome exit tunnel responsible for Trp inhibition of ribosome function. Nucleic Acids Res. 40, 2247–2257 (2012).

16. Cruz-Vera, L. R., Rajagopal, S., Squires, C. & Yanofsky, C. Features of ribosome-peptidyl-tRNA interactions essential for tryptophan induction of tna operon expression. Mol. Cell 19, 333–343 (2005).

17. Cruz-Vera, L. R., New, A., Squires, C. & Yanofsky, C. Ribosomal features essential for tna operon induction: Tryptophan binding at the peptidyl transferase center. J. Bacteriol. 189, 3140–3146 (2007).

18. Yang, R., Cruz-Vera, L. R. & Yanofsky, C. 23S rRNA nucleotides in the peptidyl transferase center are essential for tryptophanase operon induction. J. Bacteriol. 191, 3445–3450 (2009).

19. Seidelt, B. et al. Structural insight into nascent polypeptide chain–mediated translational stalling. Science 1621, 1412–1415 (2009).

20. Bischoff, L., Berninghausen, O. & Beckmann, R. Molecular basis for the ribosome functioning as an L-tryptophan sensor. Cell Rep. 9, 469–475 (2014).

21. Tian, P. et al. Folding pathway of an Ig domain is conserved on and off the ribosome. Proc. Natl. Acad. Sci. U. S. A. 115, E11284–E11293 (2018).

22. Cruz-Vera, L. R. & Yanofsky, C. Conserved residues Asp16 and Pro24 of TnaC-tRNAPro participate in tryptophan induction of tna operon expression. J. Bacteriol. 190, 4791– 4797 (2008).

23. Cruz-Vera, L. R., Yang, R. & Yanofsky, C. Tryptophan inhibits Proteus vulgaris TnaC leader peptide elongation, activating tna operon expression. J. Bacteriol. 191, 7001– 7006 (2009).

24. Pierson, W. E. et al. Uniformity of Peptide Release Is Maintained by Methylation of Release Factors. Cell Rep. 17, 11–18 (2016).

25. Martínez, A. K. et al. Interactions of the TnaC nascent peptide with rRNA in the exit tunnel enable the ribosome to respond to free tryptophan. Nucleic Acids Res. 42, 1245– 1256 (2014).

26. Herrero Del Valle, A. et al. Ornithine capture by a translating ribosome controls bacterial polyamine synthesis. Nat. Microbiol. 5, 554–561 (2020).

27. Su, T. et al. The force-sensing peptide VemP employs extreme compaction and secondary structure formation to induce ribosomal stalling. Elife 6, 1–17 (2017).

28. Gong, F., Ito, K., Nakamura, Y. & Yanofsky, C. The mechanism of tryptophan induction of tryptophanase operon expression: Tryptophan inhibits release factor-mediated cleavage of TnaC-peptidyl-tRNAPro. Proc. Natl. Acad. Sci. U. S. A. 98, 8997–9001 (2001).

29. Cruz-Vera, L. R., Gong, M. & Yanofsky, C. Changes produced by bound tryptophan in the ribosome peptidyl transferase center in response to TnaC, a nascent leader peptide. Proc. Natl. Acad. Sci. U. S. A. 103, 3598–3603 (2006).

30. Dunkle, J. A., Xiong, L., Mankin, A. S. & Cate, J. H. D. Structures of the Escherichia coli ribosome with antibiotics bound near the peptidyl transferase center explain spectra of drug action. Proc. Natl. Acad. Sci. U. S. A. 107, 17152–17157 (2010).

31. Svetlov, M. S., Vázquez-Laslop, N. & Mankin, A. S. Kinetics of drug-ribosome interactions defines the cidality of macrolide antibiotics. Proc. Natl. Acad. Sci. U. S. A. 114, 13673–13678 (2017).

32. Burakovsky, D. E. et al. Mutations at the accommodation gate of the ribosome impair RF2-dependent translation termination. Rna 16, 1848–1853 (2010).

33. Emmanuel, J. S., Sengupta, A., Gordon, E. R., Noble, J. T. & Cruz-Vera, L. R. The regulatory TnaC nascent peptide preferentially inhibits release factor 2-mediated hydrolysis of peptidyl-tRNA. J. Biol. Chem. 294, 19224–19235 (2019).

34. Uno, M., Ito, K. & Nakamura, Y. Polypeptide release at sense and noncognate stop codons by localized charge-exchange alterations in translational release factors. Proc. Natl. Acad. Sci. U. S. A. 99, 1819–1824 (2002).

35. Shaw, J. J. & Green, R. Two distinct components of release factor function uncovered by nucleophile partitioning analysis. Mol. Cell 28, 458–467 (2007).

36. Aleksashin, N. A. et al. A fully orthogonal system for protein synthesis in bacterial cells. Nat. Commun. 11, 1858 (2020).

37. Mottagui-Tabar, S., Bjornsson, A. & Isaksson, L. A. The second to last amino acid in the nascent peptide as a codon context determinant. EMBO J. 13, 249–258 (1994).

38. Kubelka, J., Hofrichter, J. & Eaton, W. A. The protein folding ‘speed limit’. Curr. Opin. Struct. Biol. 14, 76–88 (2004).

39. Li, W. et al. Structural basis for selective stalling of human ribosome nascent chain complexes by a drug-like molecule. Nat. Struct. Mol. Biol. 26, 501–509 (2019).

40. Li, W., Chang, S. T.-L., Ward, F. R. & Cate, J. H. D. Selective inhibition of human translation termination by a drug-like compound. Nat. Commun. 11, 4941 (2020).

41. Arenz, S. et al. Drug sensing by the ribosome induces translational arrest via active site perturbation. Mol. Cell 56, 446–452 (2014).

42. Arenz, S. et al. A combined cryo-EM and molecular dynamics approach reveals the mechanism of ErmBL-mediated translation arrest. Nat. Commun. 7, (2016).

43. Zhang, J. et al. Mechanisms of ribosome stalling by SecM at multiple elongation steps. Elife 4, 1–25 (2015).

44. Sohmen, D. et al. Structure of the Bacillus subtilis 70S ribosome reveals the basis for species-specific stalling. Nat. Commun. 6, 6941 (2015).

45. Dever, T. E., Ivanov, I. P. & Sachs, M. S. Conserved Upstream Open Reading Frame Nascent Peptides That Control Translation. Annu. Rev. Genet. 54, 237–264 (2020).

46. Wei, J., Wu, C. & Sachs, M. S. The arginine attenuator peptide interferes with the ribosome peptidyl transferase center. Mol. Cell. Biol. 32, 2396–2406 (2012).

47. Yamashita, Y. et al. Sucrose sensing through nascent peptide-meditated ribosome stalling at the stop codon of Arabidopsis bZIP11 uORF2. FEBS Lett. 591, 1266–1277 (2017).

48. Storz, G., Wolf, Y. I. & Ramamurthi, K. S. Small proteins can no longer be ignored. Annu. Rev. Biochem. 83, 753–777 (2014).

49. Schindelin, J. et al. Fiji: an open-source platform for biological-image analysis. Nat. Methods 9, 676–682 (2012).

50. Mastronarde, D. N. Automated electron microscope tomography using robust prediction of specimen movements. J. Struct. Biol. 152, 36–51 (2005).

51. Zivanov, J., Nakane, T. & Scheres, S. H. W. Estimation of high-order aberrations and anisotropic magnification from cryo-EM data sets in RELION-3.1. IUCrJ 7, 253–267 (2020).

52. Zheng, S. Q. et al. MotionCor2: anisotropic correction of beam-induced motion for improved cryo-electron microscopy. Nat. Methods 14, 331–332 (2017).

53. Zhang, K. Gctf: Real-time CTF determination and correction. J. Struct. Biol. 193, 1–12 (2016).

54. Wilkinson, M. E., Kumar, A. & Casañal, A. Methods for merging data sets in electron cryo-microscopy. Acta Crystallogr. Sect. D, Struct. Biol. 75, 782–791 (2019).

55. Terwilliger, T. C., Sobolev, O. V, Afonine, P. V & Adams, P. D. Automated map sharpening by maximization of detail and connectivity. Acta Crystallogr. Sect. D, Struct. Biol. 74, 545–559 (2018).

56. Pettersen, E. F. et al. UCSF ChimeraX: Structure visualization for researchers, educators, and developers. Protein Sci. 30, 70–82 (2021).

57. Emsley, P., Lohkamp, B., Scott, W. G. & Cowtan, K. Features and development of Coot. Acta Crystallogr. D. Biol. Crystallogr. 66, 486–501 (2010).

58. Afonine, P. V et al. Real-space refinement in PHENIX for cryo-EM and crystallography. Acta Crystallogr. Sect. D, Struct. Biol. 74, 531–544 (2018).

59. Croll, T. I. ISOLDE: a physically realistic environment for model building into low-resolution electron-density maps. Acta Crystallogr. Sect. D, Struct. Biol. 74, 519–530 (2018).

60. Fu, Z. et al. The structural basis for release-factor activation during translation termination revealed by time-resolved cryogenic electron microscopy. Nat. Commun. 10, 1–7 (2019).

61. Liebschner, D. et al. Macromolecular structure determination using X-rays, neutrons and electrons: recent developments in Phenix. Acta Crystallogr. Sect. D, Struct. Biol. 75, 861–877 (2019).

62. Kandiah, E. et al. CM01: A facility for cryo-electron microscopy at the European synchrotron. Acta Crystallogr. Sect. D Struct. Biol. 75, 528–535 (2019).

63. Korostelev, A. et al. Crystal structure of a translation termination complex formed with release factor RF2. Proc. Natl. Acad. Sci. U. S. A. 105, 19684–19689 (2008).

